# Anti-chemokine antibodies after SARS-CoV-2 infection correlate with favorable disease course

**DOI:** 10.1101/2022.05.23.493121

**Authors:** Jonathan Muri, Valentina Cecchinato, Andrea Cavalli, Akanksha A. Shanbhag, Milos Matkovic, Maira Biggiogero, Pier Andrea Maida, Jacques Moritz, Chiara Toscano, Elaheh Ghovehoud, Raffaello Furlan, Franca Barbic, Antonio Voza, Guendalina De Nadai, Carlo Cervia, Yves Zurbuchen, Patrick Taeschler, Lilly A. Murray, Gabriela Danelon-Sargenti, Simone Moro, Tao Gong, Pietro Piffaretti, Filippo Bianchini, Virginia Crivelli, Lucie Podešvová, Mattia Pedotti, David Jarrossay, Jacopo Sgrignani, Sylvia Thelen, Mario Uhr, Enos Bernasconi, Andri Rauch, Antonio Manzo, Adrian Ciurea, Marco B.L. Rocchi, Luca Varani, Bernhard Moser, Barbara Bottazzi, Marcus Thelen, Brian A. Fallon, Onur Boyman, Alberto Mantovani, Christian Garzoni, Alessandra Franzetti-Pellanda, Mariagrazia Uguccioni, Davide F. Robbiani

## Abstract

Infection by SARS-CoV-2 leads to diverse symptoms, which can persist for months. While antiviral antibodies are protective, those targeting interferons and other immune factors are associated with adverse COVID-19 outcomes. Instead, we discovered that antibodies against specific chemokines are omnipresent after COVID-19, associated with favorable disease, and predictive of lack of long COVID symptoms at one year post infection. Anti-chemokine antibodies are present also in HIV-1 infection and autoimmune disorders, but they target different chemokines than those in COVID-19. Monoclonal antibodies derived from COVID- 19 convalescents that bind to the chemokine N-loop impair cell migration. Given the role of chemokines in orchestrating immune cell trafficking, naturally arising anti-chemokine antibodies associated with favorable COVID-19 may be beneficial by modulating the inflammatory response and thus bear therapeutic potential.

**One-Sentence Summary:** Naturally arising anti-chemokine antibodies associate with favorable COVID-19 and predict lack of long COVID.

---

The spectrum of disease manifestations upon infection with severe acute respiratory syndrome coronavirus 2 (SARS-CoV-2) is broad ^1^. Factors that predispose people to hospitalization and death include age, gender, ethnicity, obesity, genetic predisposition, autoantibodies against interferon, and comorbidities such as hypertension, diabetes, and coronary heart disease ^2–6^. Coronavirus disease 2019 (COVID-19) convalescent individuals often lament protracted symptoms over months, a condition referred to as long COVID or PASC (Post-Acute Sequelae of COVID), and are at increased risk of cardiovascular events ^7–13^. Some evidence points to a role for immune dysregulation and autoimmunity as contributors to long COVID, although virus persistence has also been proposed ^14–16^. Overall, there is however little understanding of the biology underlying long COVID and of the reasons for the differences in COVID-19 manifestation.

Chemokines are chemotactic cytokines that mediate leukocyte trafficking and activity by binding to seven-transmembrane G protein-coupled receptors ^17, 18^. They play a fundamental role in health and disease since the proper trafficking of leukocyte subsets is governed by the combinatorial diversity of their responsiveness to chemokines ^18^. In addition to elevated levels of pro-inflammatory cytokines (*e.g.* IL-6, TNF, and IL1β), higher levels of certain chemokines are observed in acute COVID-19 (*e.g.* CCL2, CCL3, CCL4, CCL7, CCL8, CCL19, CXCL2, CXCL5, CXCL8, CXCL9, CXCL10, CXCL13, CXCL16 and CXCL17) and multiomic studies recently identified plasma chemokines among the most significant factors associated with COVID-19 severity ^19–26^. Accordingly, chemokines recruit neutrophils and monocytes to sites of infection, where they play a key role in the pathophysiology of COVID-19 by sustaining inflammation and causing collateral tissue damage and fibrosis, particularly during the inflammatory phase that follows virus clearance ^20, 24, 27–30^. Chemokines have also been implicated in the pathogenesis and as biomarkers of long COVID ^31^. Anti-inflammatory treatments, such as steroids and IL-6 blockade, are efficacious in hospitalized COVID-19 patients, and therapies targeting the chemokine system are under development for immunological disorders and have been proposed for COVID-19 ^18, 22, 32, 33^.

Similar to earlier work linking anti-cytokine antibodies to mycobacterial, staphylococcal and fungal diseases ^34–36^, autoantibodies against cytokines have been described in COVID-19. In particular, anti-type I Interferon antibodies distinguished ∼10% of life-threatening pneumonia and ∼20% of deaths from COVID-19 ^6, 37, 38^. Moreover, autoantibodies characteristic of systemic autoimmune disorders, such as anti-phospholipid antibodies, anti-nuclear antibodies and rheumatoid factor, were reported in COVID-19 ^39–43^. A recent high-throughput screening by yeast display of the secretome further revealed the presence of autoantibodies against several immune factors, including chemokines ^44^. However, anti-chemokine antibodies were infrequent by this method, and there was neither correlation with disease severity, or long COVID, nor information about the persistence of such autoantibodies over time.

We devised a peptide-based strategy to measure and discover antibodies that bind to a functional region of each of the 43 human chemokines. By examining three diverse independent cohorts, comprising subjects who contracted COVID-19 during 2020 and early 2021, we found that the presence of antibodies against specific chemokines helps to identify convalescent individuals with favorable acute and long COVID disease course. Anti- chemokine monoclonal antibodies derived from these individuals block leukocyte migration and thus may be beneficial through modulation of the inflammatory response.

## RESULTS

### COVID-19 and anti-chemokine antibodies

To evaluate anti-chemokine antibodies after COVID-19, we obtained plasma samples from a convalescent cohort on average at 6 months after disease onset (Lugano cohort; Extended Data Fig. 1a; Supplementary Table 1). Since the N-terminal loop (N-loop) of chemokines is required for receptor binding, we reasoned that biologically active anti-chemokine antibodies would likely target this region, whose sequence happens to be specific for each but two of the 43 human chemokines (Extended Data Fig. 1b) ^45^. Therefore, we designed peptides corresponding to the N-loop of each chemokine for use in enzyme-linked immunosorbent assays (ELISA, Supplementary Table 2). IgG antibody levels were measured on serial plasma dilutions and the signal plotted as heatmap (Fig. 1a; Extended Data Fig. 1c; Supplementary Table 3). Analysis of all parameters by nonlinear dimensionality reduction with *t*-distributed stochastic neighbor embedding (t-SNE) revealed a clear separation between controls and COVID-19 convalescents (Extended Data Fig. 2a). Some convalescent plasma revealed high levels of IgGs to certain chemokines (for example CCL8, CXCL13 and CXCL16). For these chemokines, antibody levels to the N-loop significantly correlated with those against the C-terminal region of the same chemokine, suggesting that, when present, antibodies formed against multiple chemokine epitopes (Fig. 1a; Extended Data Fig. 2b). When considering antibodies against each chemokine individually, a significant difference in reactivity over uninfected controls was observed for peptides corresponding to 23 of the 43 chemokines (Extended Data Fig. 2c). Antibodies to the three chemokines with p<10^-4^ (CCL19, CCL22 and CXCL17; “COVID-19 signature”) clustered together, and by themselves were sufficient to correctly assign uninfected controls and COVID-19 convalescents with high accuracy (96.8%; Fig. 1a-c; Extended Data Fig. 2c,d; see Methods). The findings were validated with two independent cohorts: one that was sampled during the acute phase and on average at 7 months from disease onset (Milan cohort, n=44; 90.5% and 89.5% accuracy, respectively; Extended Data Fig. 3a,b; Supplementary Tables 1 and 3), and a second cohort that was evaluated at 13 months (Zurich cohort, n=104; 92.9% accuracy; Extended Data Fig. 3a,b; Supplementary Tables 1 and 3). Thus, COVID-19 is associated with a specific pattern of anti-chemokine antibodies.

**Fig. 1.**
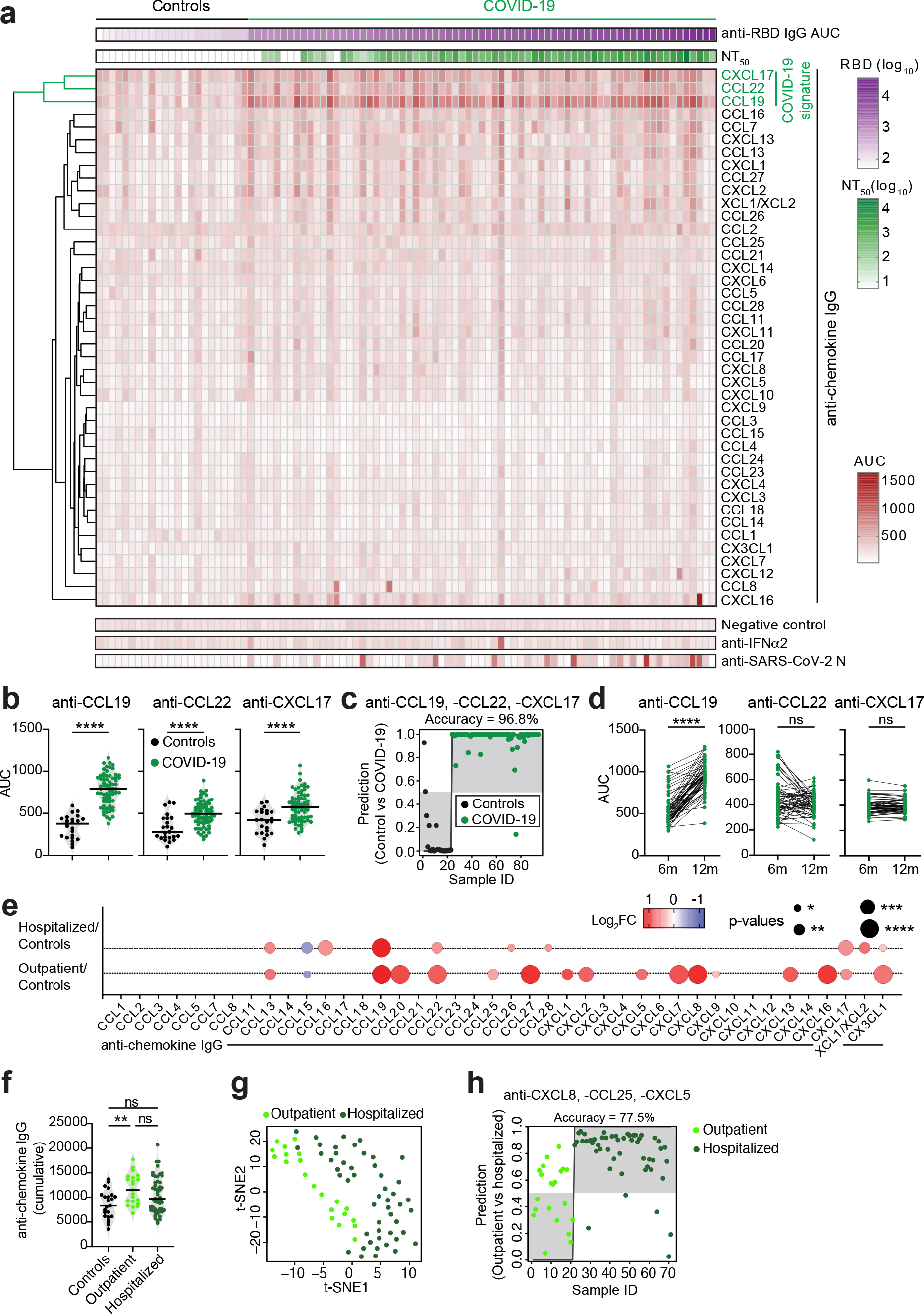
Distinct patterns of anti-chemokine antibodies in COVID-19 convalescents with different severity of acute disease. (**a**) Anti-chemokine antibodies 6 months after COVID-19. Heatmap representing plasma IgG binding to 42 peptides comprising the N-loop sequence of all 43 human chemokines, as determined by ELISA (Area Under the Curve [AUC], average of two independent experiments). Samples are ranked according to the level of anti-SARS-CoV- 2-RBD reactivity. Anti-chemokine IgGs are ordered by unsupervised clustering analysis of ELISA signal. SARS-CoV-2 pseudovirus neutralizing activity (NT50) and IgG binding to peptides corresponding to negative control, IFNα2 and SARS-CoV-2 nucleocapsid protein (N) are shown. COVID-19 convalescents (n=71); controls (n=23). (**b**) Difference in IgG antibodies to CCL19, CCL22 and CXCL17 (COVID-19 signature). Horizontal bars indicate median values. Two-tailed Mann–Whitney U-tests. (**c**) Assignment of COVID-19 convalescents and controls based on the COVID-19 signature antibodies by logistic regression analysis. Dots on grey background are correctly assigned. (**d**) Anti-COVID-19 signature chemokine IgG antibodies at 6 and 12 months in convalescents. AUC from two independent experiments. Wilcoxon signed-rank test. (**e**) Difference in anti-chemokine antibodies between COVID-19 groups and controls. Summary circle plot: circle size indicates significance; colors show the Log2 fold-change increase (red) or decrease (blue) over controls. Kruskal-Wallis test followed by Dunn’s multiple comparison test. (**f**) Difference in total anti-chemokine antibodies. Cumulative signal of the IgGs against the 42 peptides comprising the N-loop sequence of all 43 human chemokines. Horizontal bars indicate median values. Kruskal-Wallis test followed by Dunn’s multiple comparison test. (**g**) t-SNE distribution of COVID-19 outpatient and hospitalized individuals, as determined with the 42 datasets combined. (**h**) Assignment of COVID-19 outpatient and hospitalized individuals based on the COVID-19 hospitalization signature antibodies by logistic regression analysis. Dots on grey background are correctly assigned. See also Extended Data Figs. 1-6.

**Fig. 2.**
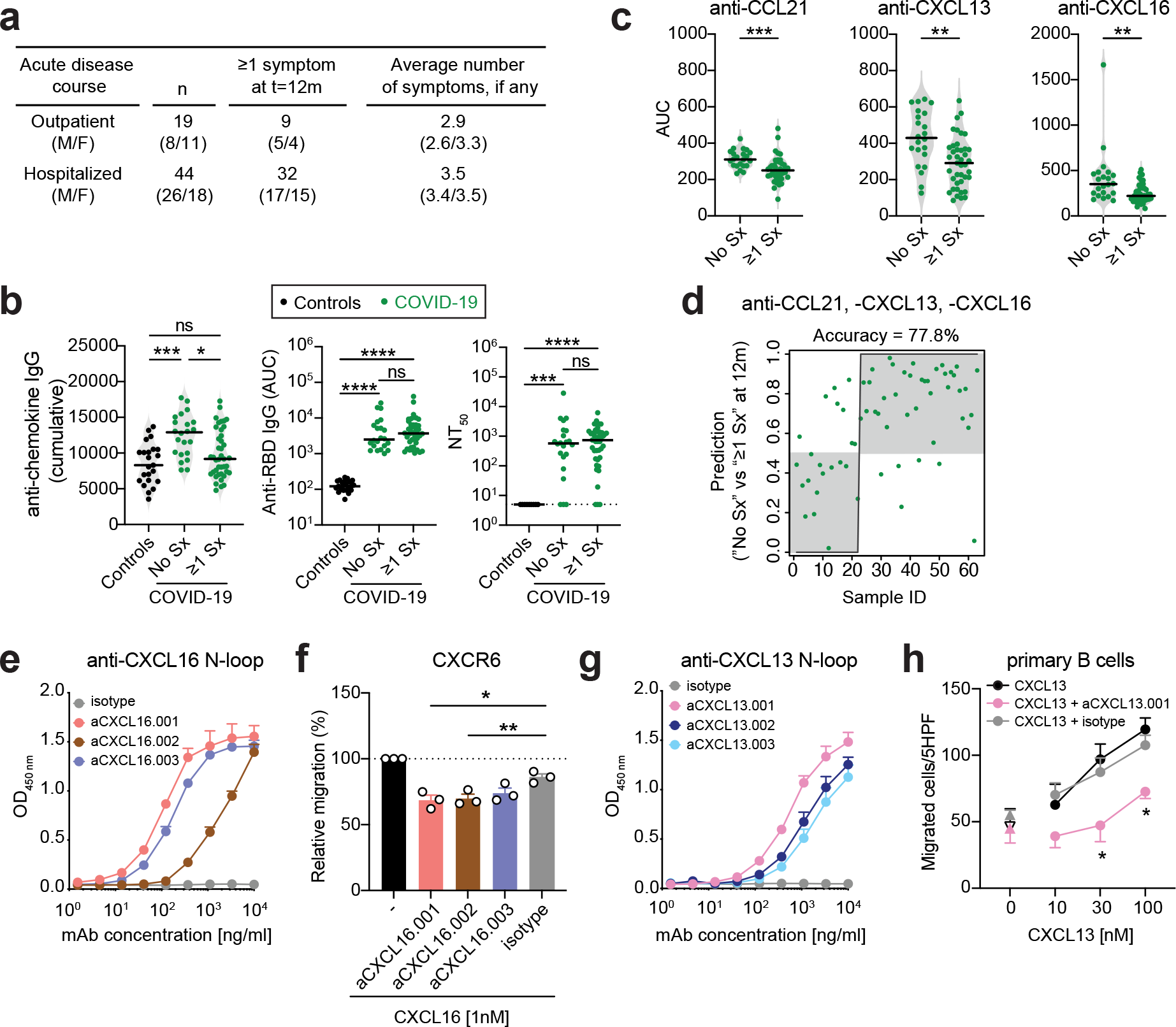
Anti-chemokine antibodies and long COVID. (**a**) Characteristics of the COVID-19 convalescent Lugano cohort at 12 months. (**b**) Persisting symptoms (Sx) at 12 months and anti-chemokine IgG (cumulative; left), anti-RBD IgG (middle), and NT50 (right) values at 6 months. Horizontal bars indicate median values. Average AUC from two independent experiments. Kruskal-Wallis test followed by Dunn’s multiple comparison test. (**c**) Difference in antibodies to CCL21, CXCL13 and CXCL16 (Long COVID signature). Horizontal bars indicate median values. Average AUC from two independent experiments. Two-tailed Mann–Whitney U-tests. (**d**) Group assignment based on the Long COVID signature antibodies at 6 months against CCL21, CXCL13 and CXCL16, by logistic regression analysis. Dots on grey background are correctly assigned. (**e**) Anti-CXCL16 antibodies binding to the CXCL16 N-loop in ELISA. Average of two independent experiments (Mean+SEM). (**f**) Inhibition of chemotaxis by anti-CXCL16 N-loop antibodies. Relative cell migration towards CXCL16 by cells uniquely expressing CXCR6 (see Methods). Mean+SEM of 3 independent experiments. Paired, two-tailed Student’s t test. (**g**) Anti-CXCL13 antibodies binding to the CXCL13 N-loop in ELISA. Average of two independent experiments (Mean+SEM). (**h**) The anti-CXCL13 N-loop antibody aCXCL13.001 inhibits CXCL13 chemotaxis of primary CD19^+^ human B cells. Mean±SEM of migrated cells in 5 high-power fields (HPF). Average of 3 independent experiments with cells from different donors. Up-pointing triangles indicate antibody alone, and down-pointing triangle is buffer control. Two-way RM ANOVA followed by Šídák’s multiple comparisons test. See also Extended Data Figs. 7 and 8.

**Fig. 3.**
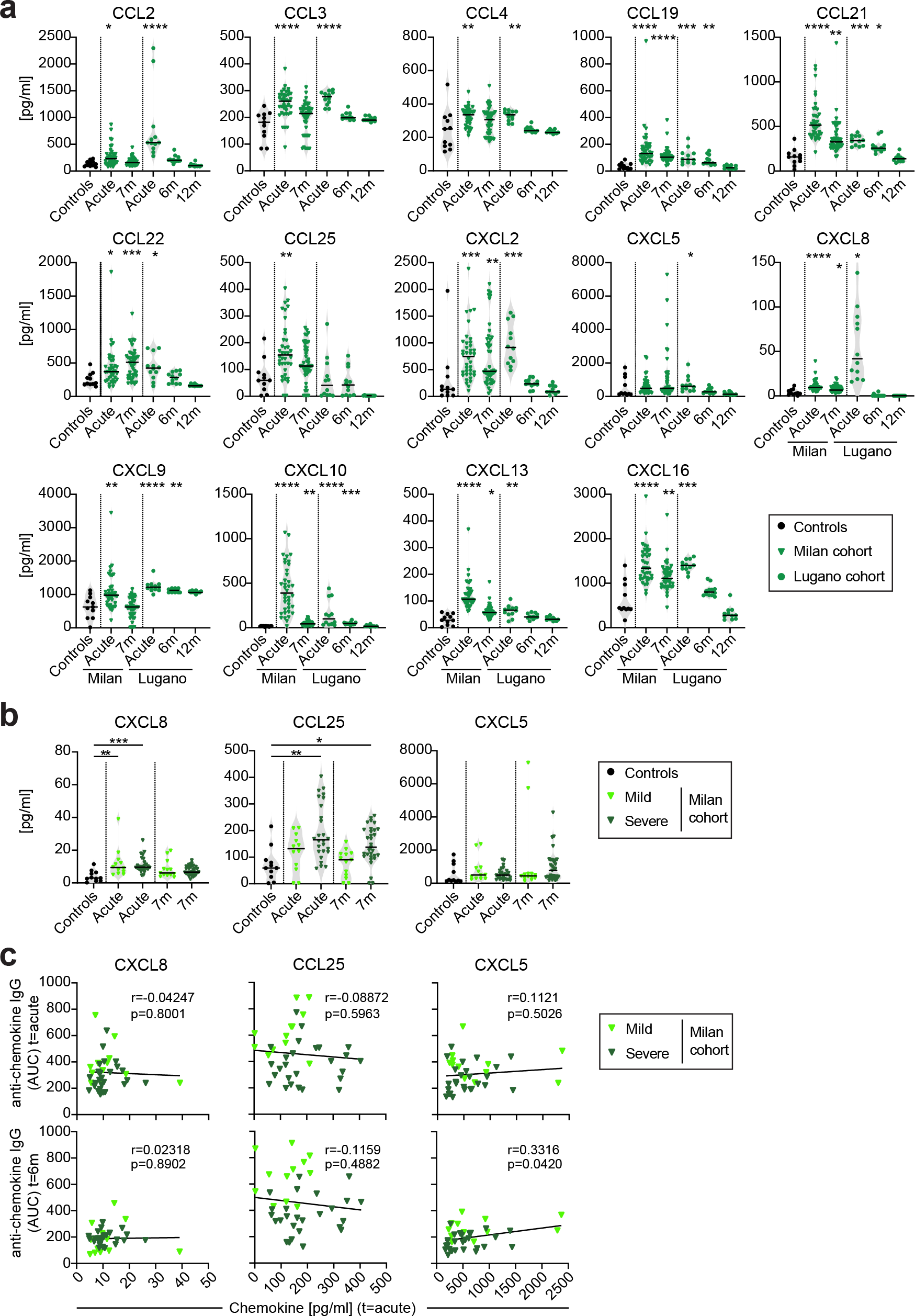
Chemokines in plasma during or after COVID-19. (**a**) Plasma chemokine levels in the Milan (n=44) and Lugano (n=12) cohorts at the indicated time points after disease onset. Horizontal bars indicate median values. Kruskal-Wallis test followed by Dunn’s multiple comparison test over controls. (**b**) Levels of COVID-19 hospitalization signature chemokines (CXCL5, CXCL8 and CCL25) in mild versus severe patients (Milan cohort). Horizontal bars indicate median values. Kruskal-Wallis test followed by Dunn’s multiple comparison test. (**c**) Correlation between chemokine and autoantibody levels determined by Pearson correlation analysis.

To examine the relationship between anti-chemokine antibodies and other serologic features of the COVID-19 cohort, we used ELISA and a pseudovirus-based neutralization assay to measure binding and neutralizing capacity of antibodies against SARS-CoV-2 (Fig. 1a) ^46^.

In agreement with previous studies, IgG binding to SARS-CoV-2 Spike receptor binding domain (RBD) and plasma half-maximal SARS-CoV-2 neutralizing titers (NT50) were variable but positively correlated with each other and with age (Extended Data Fig. 4a-c) ^46^. In contrast, there was no correlation between NT50 or anti-RBD IgGs and the levels of antibodies to the signature chemokines CCL19, CCL22 and CXCL17, or to the sum of all anti-chemokine IgG reactivities (cumulative area under the curve; Extended Data Fig. 4d). A weak negative correlation between age and the sum of all anti-chemokine IgG reactivities was observed (Extended Data Fig. 4d), but there were no differences in the levels of antibodies to the signature chemokines between males and females (Extended Data Fig. 4e). We conclude that, after COVID-19, antibodies against specific chemokines are not correlated with those against SARS-CoV-2.

**Fig. 4.**
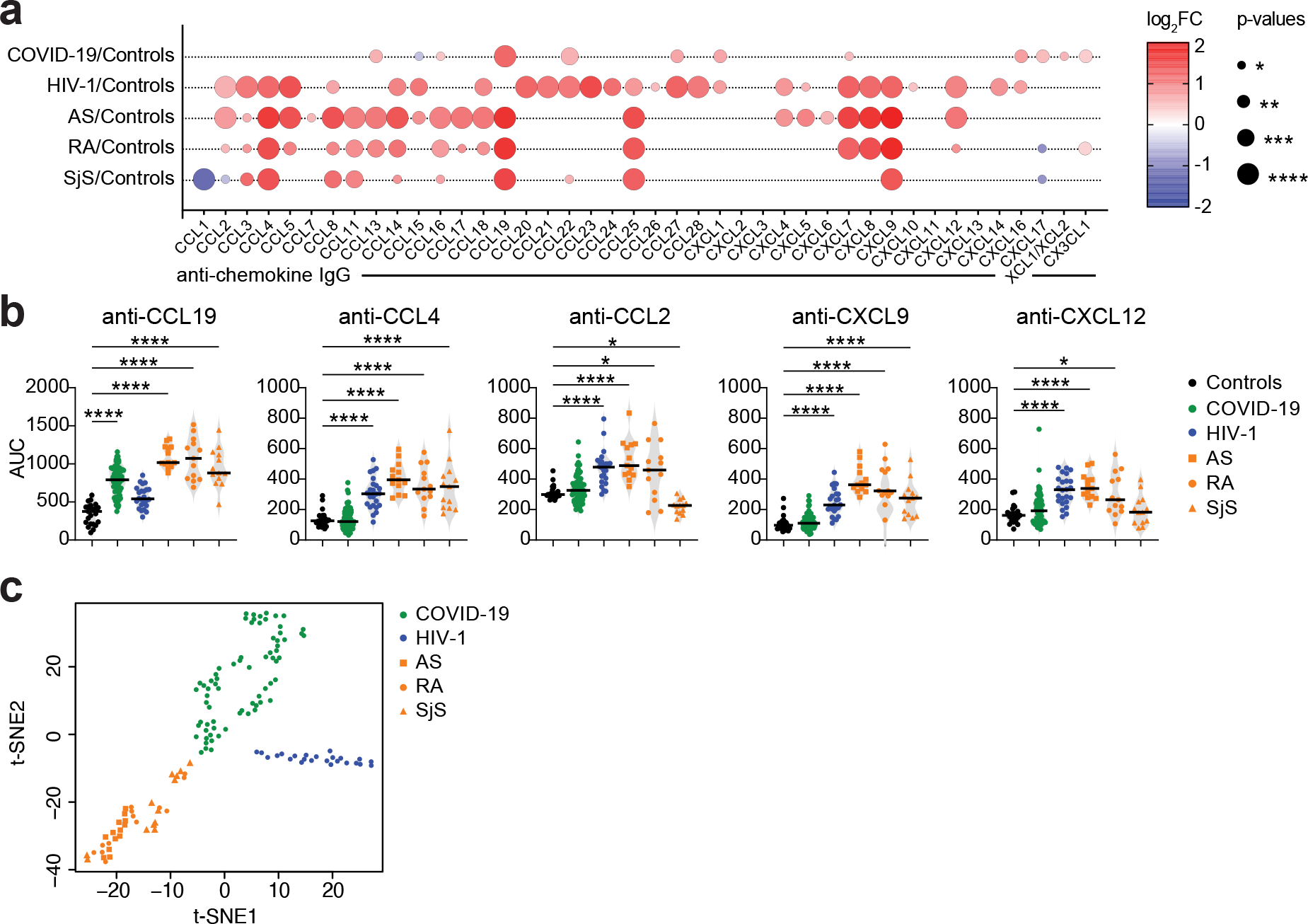
Distinct patterns of anti-chemokine antibodies in COVID-19, HIV-1 or autoimmune diseases. (**a**) Difference in anti-chemokine antibodies between diseases and controls. Summary circle plot: circle size indicates significance; colors show the Log2 fold- change increase (red) or decrease (blue) over controls. Kruskal-Wallis test followed by Dunn’s multiple comparison test. (**b**) Difference in antibodies to CCL19, CCL4, CCL2, CXCL9 and CXCL12 across groups. Controls (n=23), COVID-19 (n=71), HIV-1 (n=24), Ankylosing Spondylitis (AS, n=13), Rheumatoid Arthritis (RA, n=13), and Sjögren’s syndrome (SjS, n=13). Horizontal bars indicate median values. Average AUC from two independent experiments. Kruskal-Wallis test followed by Dunn’s multiple comparison test over rank of the control group. (**c**) t-SNE distribution of disease samples, as determined with the 42 datasets combined. See also Extended Data Figs. 9 and 10.

### Anti-chemokine antibodies over time

To document the temporal evolution of anti-chemokine antibodies following COVID-19, we compared side-by-side the reactivities of plasma collected from the Lugano cohort at approximately 6 and 12 months from symptom onset (Extended Data Fig. 5a; Supplementary Tables 1 and 3). In agreement with earlier findings ^47, 48^, antibodies to the virus RBD significantly decreased in unvaccinated COVID-19 convalescents, while they increased in those receiving at least one dose of mRNA-based COVID-19 vaccine (Extended Data Fig. 5b; Supplementary Table 1). Conversely, and regardless of vaccination status, antibodies to the COVID-19 signature chemokine CCL19 significantly increased (2.1-fold, p<0.0001), those to CXCL17 remained generally stable, and those to CCL22 followed variable kinetics (Fig. 1d and data not shown). Similar to those against CCL19, antibody levels to CCL8, CCL13, CCL16, CXCL7 and CX3CL1 were also augmented at 12 months (Extended Data Fig. 5c). To further investigate the kinetics of COVID-19 signature antibodies, we analyzed cohort individuals for which acute samples were also available (n=12; Extended Data Fig. 5d; Supplementary Tables 1 and 3). During acute COVID-19, IgG antibodies to CCL19, but not to CCL22 or CXCL17, were already higher than in uninfected controls, and continued to increase until 12 months (Extended Data Fig. 5e). Similarly, in the Milan cohort, autoantibodies against these three chemokines were already higher than in controls during the acute phase (Extended Data Fig. 3b; Supplementary Table 3). In contrast to natural infection, no significant change in antibody reactivity to any of the chemokines was observed upon COVID-19 mRNA vaccination of SARS-CoV-2 naïve individuals after about 4 months (130 days on average; n=16; Extended Data Fig. 5f; Supplementary Tables 1 and 3). Therefore, unlike the antibodies to SARS-CoV-2 RBD, which decrease over time, the levels of some anti-chemokine antibodies that are present upon COVID-19 increase over one year of observation.

### Anti-chemokine antibodies and severity of disease

Autoantibodies have been detected in a portion of hospitalized COVID-19 patients, linking their presence to severe illness ^6, 39, 43, 44^. To evaluate the relationship between the severity of acute COVID-19 and convalescent anti-chemokine IgGs, we compared individuals in the original Lugano cohort who were either hospitalized because of the infection (n=50) or remained as outpatients (n=21; Fig. 1e). No significant difference in age distribution was observed between groups (age [years]: mean±SD; 60±14 in hospitalized, 57±15 in outpatients; p=0.3487), while a higher proportion of males was observed among hospitalized but not outpatients (60% and 38.1%, respectively; Supplementary Table 1) ^5^.

When the most significant differences in autoantibody levels were considered (p<10^-4^), only the antibodies against CCL19 were higher in hospitalized individuals over uninfected controls, while antibodies against 8 chemokines (CXCL8, CCL22, CXCL16, CCL27, CXCL7, CCL20, CX3CL1, in addition to CCL19) were increased in outpatients (Fig. 1e; Extended Data Fig. 6a). Consistent with this finding, the outpatient but not the hospitalized individuals displayed significantly higher cumulative anti-chemokine reactivity (p=0.0038; Fig. 1f). Thus, a broader pattern and higher overall amounts of anti-chemokine antibodies are observed at 6 months in those COVID-19 convalescents, who were outpatients during the acute phase of the disease.

Direct comparison of previously hospitalized and outpatient individuals by t-SNE analysis of all anti-chemokine datasets separated the two groups (Fig. 1g). Antibodies against three chemokines highly significantly distinguished outpatients from hospitalized subjects (p<10^-4^): antibodies against CXCL5, CXCL8 and CCL25 were all lower in individuals with severe illness requiring hospitalization, and this was not linked to the therapy received during hospitalization (Extended Data Fig. 6a,b; Supplementary Table 1). The combination of antibody values against these three chemokines alone could correctly assign formerly hospitalized and outpatient individuals with an accuracy of 77.5% (“COVID-19 hospitalization signature”; Fig. 1h). Similar findings were obtained with the Milan (85.0% and 84.1% accuracy at acute and 7 months, respectively; Extended Data Fig. 3c) and Zurich cohorts (73.1% accuracy at 13 months; Extended Data Fig. 3c). Among the COVID-19 hospitalization signature antibodies, those to CXCL5 and CXCL8 were negatively correlated with anti-RBD IgG and age, while no differences were observed between males and females (Extended Data Fig. 6c,d). Consistent with previous work, both anti-RBD IgG and NT50 values were significantly higher in hospitalized individuals compared to outpatients and in males compared to females of both groups (Extended Data Fig. 6e) ^46^. Thus, the anti-chemokine antibody signature that distinguishes uninfected from COVID-19 convalescents (CCL19, CCL22 and CXCL17; Fig. 1b,c) is different from the signature associated with different severity of COVID-19 disease (CXCL5, CXCL8 and CCL25; Fig. 1h).

### Anti-chemokine antibodies and long COVID

A fraction of individuals who recover from COVID-19 experience long-term sequelae ^7–10^. To determine whether a specific pattern of anti-chemokine antibodies at 6 months is predictive of the persistence of symptoms, we collected information on long COVID from the Lugano cohort at 12 months (Fig. 2). 65.1% of all participants reported persistence of at least one symptom related to COVID-19. Among these, the average number of long-term symptoms was 3.3, and they were more frequent among formerly hospitalized individuals than outpatients (72.7% versus 47.4%; Fig. 2a; Extended Data Fig. 7a,b; Supplementary Table 1). No differences in age, gender distribution or time from disease onset to second visit were observed between individuals with and without protracted symptoms (Extended Data Fig. 7c).

Convalescents with long-term sequelae showed significantly lower cumulative levels of anti-chemokine antibodies compared to those without symptoms (p=0.0135; Fig. 2b). This was particularly true for outpatients and among females (Extended Data Fig. 7d,e). In contrast, anti-RBD IgG and NT50 values were comparable between the two groups (Fig. 2b). The total levels of anti-chemokine antibodies did not correlate with the number of symptoms (Extended Data Fig. 7f). These data indicate that overall higher levels of anti-chemokine antibodies at 6 months after COVID-19 are associated with absence of long-term symptoms at 12 months.

IgG antibodies against three chemokines distinguished the groups with high significance: CCL21 (p=0.0001), CXCL13 (p=0.0010) and CXCL16 (p=0.0011; Fig. 2c; Extended Data Fig. 7g; “Long COVID signature”). Logistic regression analysis using the antibody values for these 3 chemokines alone predicted the absence of persistent symptoms with 77.8% accuracy (Fig. 2d). Similarly, analysis of the Zurich cohort at 13 months showed 72.1% accuracy of association with lack of long COVID, even though in that cohort only anti-CCL21 antibodies were significantly different between groups (Extended Data Fig. 3d). These results indicate that specific patterns of anti-chemokine antibodies at 6 months are associated with the longer-term persistence of symptoms after COVID-19.

Since anti-chemokine antibodies to CXCL13 and CXCL16 are associated with decreased likelihood of long COVID, we next derived corresponding memory B cell antibodies from available PBMC samples (Supplementary Table 4; see Methods). Three N-loop binding monoclonal antibodies were obtained for CXCL16, which blocked migration of a cell line expressing the cognate receptor (CXCR6; Fig. 2e,f; Extended Data Fig. 8a,b; Supplementary Tables 4 and 5). Similarly, 3 anti-CXCL13 N-loop antibodies bound in ELISA and inhibited chemotaxis of primary CD19^+^ human B cells (Fig. 2g,h; Extended Data Fig. 8c; Supplementary Tables 4 and 5). By the same approach, we discovered chemotaxis-blocking antibodies specific for CCL8 and CCL20 (Extended Data Fig. 8d-j and Supplementary Tables 4 and 5). Consistent with these results, polyclonal plasma IgG from COVID-19 convalescents effectively blocked chemotaxis at concentrations 50 times lower than those found in human serum^49^ (Extended Data Fig. 8k). Therefore, confirming our initial hypothesis, antibodies from COVID-19 convalescents that bind to the N-loop of chemokines are biologically active.

### Lack of correlation between chemokines and their antibodies

To test the correlation between autoantibody and corresponding antigen, we first measured plasma chemokine levels. In agreement with earlier reports^19–25^, the levels of CCL2, CCL3, CCL4, CCL19, CCL21, CCL22, CCL25, CXCL2, CXCL8, CXCL9, CXCL10, CXCL13 and CXCL16 were significantly elevated during acute disease, and 8 of these chemokines remained above control levels at 7 months post infection (CCL19, CCL21, CCL22, CXCL2, CXCL8, CXCL10, CXCL13 and CXCL16; Milan cohort; Fig. 3a and Supplementary Table 6). Similar results were observed with the 12 individuals from the Lugano cohort for which acute samples were available (Fig. 3a). Of note, none of the chemokines corresponding to autoantibodies of the hospitalization signature (CXCL5, CXCL8 and CCL25) were significantly different between mild and severe patients (Milan cohort; Fig. 3b).

We next compared the levels of chemokine and corresponding autoantibody. Consistent with the lack of significant differences in chemokine levels between groups, no correlation was observed between the levels of chemokines and those of the related signature autoantibodies in the acute phase or at 7 months post infection (Milan cohort; Fig. 3c; Supplementary Table 6). Furthermore, no increase of anti-CCL3 and anti-CCL4 antibodies was detected, although the amount of corresponding chemokines was elevated in plasma (Lugano cohort; Fig. 3b and Extended Data Fig. 6a). We conclude that, even though chemokines rapidly increase and persist for at least 6 months in plasma, their level does not correlate with the amount of corresponding anti-chemokine antibodies in the circulation.

### Anti-chemokine antibodies in other infectious and autoimmune diseases

To examine the relevance of anti-chemokine antibodies beyond COVID-19, we measured their presence in plasma from patients chronically infected with HIV-1 (n=24), and from individuals affected by Ankylosing Spondylitis (AS, n=13), Rheumatoid Arthritis (RA, n=13) and Sjögren Syndrome (SjS, n=13; Fig. 4; Supplementary Tables 1 and 3; see Methods). While antibodies against a single chemokine distinguished COVID-19 from uninfected controls with high confidence (CCL19, p<10^-4^), in HIV-1, antibodies against 14 chemokines (that did not include CCL19) were significantly increased: CCL2, CCL3, CCL4, CCL5, CCL20, CCL21, CCL22, CCL23, CCL27, CCL28, CXCL7, CXCL8, CXCL9 and CXCL12 (p<10^-4^ for all; Fig. 4a,b; Extended Data Fig. 9a; Supplementary Table 3). Similarly, AS, RA and SjS shared autoantibodies against 4 chemokines: CCL4, CCL19, CCL25 and CXCL9 (p<10^-4^ for all; Fig. 4a,b; Extended Data Fig. 9a and Extended Data Fig. 10a). In contrast, samples from *Borrelia* infected individuals (Lyme disease cohort; n=27) were indistinguishable from control except for elevated anti-CXCL14 antibodies in the acute phase (Extended Data Fig. 9b, Supplementary Tables 1 and 3). Unsupervised clustering analysis with all anti-chemokine antibody values correctly categorized all COVID-19 and HIV-1 samples with 100% accuracy, while the autoimmune diseases all clustered with each other (Extended Data Fig. 10b; see Methods). A similar result was obtained by t-SNE analysis (Fig. 4c). Thus, patterns of anti-chemokine antibodies not only distinguish different COVID-19 trajectories, but also characterize other infections and autoimmune disorders.

## DISCUSSION

We discovered that autoantibodies against chemokines are omnipresent after SARS-CoV-2 infection, and that higher levels of specific anti-chemokine antibodies are associated with favorable disease outcomes. Our findings, validated by two additional cohorts, contrast previous reports that connected autoantibodies to severe COVID-19 illness. For example, autoantibodies against type I interferon were detected in 10-20% of individuals with COVID-19 pneumonia or dying from COVID-19 ^6, 37^, and autoantibodies against a panel of immune molecules (including chemokines) and other self-antigens were described to occur sporadically and more frequently in critical COVID-19 ^39, 40, 44, 50^. These observations are in line with earlier work linking the presence of autoantibodies to adverse outcome in other infections^34–36, 51, 52^.

Chemokines drive the activation and recruitment of leukocytes to sites of infection and are involved in tissue repair ^17, 18^. Accordingly, in COVID-19, several chemokines are detected in high amounts in bronchoalveolar and other fluids, fueling a pro-inflammatory environment in the lungs, which likely contributes to COVID-19 critical illness and hospitalization ^19–23^. We find the levels of autoantibodies against CXCL5, CXCL8 and CCL25 to be augmented in COVID-19 patients with milder disease over those that require hospitalization already during the acute phase. Interestingly, the plasma levels of all three corresponding chemokines were also increased during acute disease, but they could not distinguish the two groups and there was no correlation between levels of chemokine and corresponding autoantibody. Since these chemokines attract neutrophils and other cell types that promote inflammation and tissue remodeling, the presence of the corresponding autoantibodies suggests protection through dampening of the damaging inflammatory response associated with severe COVID-19. A disease-modifying role of the chemokine system in acute COVID-19 is further supported by transcriptomic analyses and by genetic studies identifying regions of chromosome 3 encoding for chemokine receptors to be linked to critical illness ^3, 53, 54^.

Like those associated with milder disease, autoantibodies to three other chemokines (CCL21, CXCL13 and CXCL16) are increased in individuals without long COVID one year after the infection. These chemokines are important for tissue trafficking and activation of T and B lymphocytes. Therefore, it is conceivable that the corresponding autoantibodies positively impact the long-term outcome of COVID-19 by antagonizing or otherwise modulating the activation, recruitment and retention of these cell types ^55^. In keeping with this observation, persistent immune responses have been proposed as a mechanism for long COVID, and chemokines have been implicated in its pathogenesis ^7, 31^.

Regardless of disease trajectory, the presence of three other anti-chemokine antibodies is generally associated with COVID-19 infection: CCL19, CCL22 and CXCL17. Interestingly, IgG autoantibodies to these three chemokines are detected early on during the acute phase, suggesting that they are either pre-existing or rapidly induced following the infection. The early detection of anti-chemokine antibodies and their persistence or even increase between 6 and 12 months from disease onset is consistent with the rapid upregulation of the corresponding chemokines during COVID-19 and the observation that their plasma concentration remains above baseline for up to at least 6 months post infection. This is unlikely to be related to chronic SARS-CoV-2 infection because antiviral antibodies decrease during this time ^47^. Rather, the findings are consistent with the persistence of the autoantigen within germinal centers, leading to continuous generation of antibody-secreting plasma cells ^56^. We cannot exclude that the overall lack of correlation between plasma chemokine concentration and autoantibody level may be due to timing of sampling and different half-life of antibodies and chemokines in plasma, or that plasma levels may not reflect chemokine concentrations in tissues that could be more relevant for antibody induction. Further highlighting the complexity of the phenomenon, antibodies are not induced against some of the chemokines that are remarkably increased during COVID-19 (*e.g.* CCL3, CCL4 and CXCL9).

Chemokines play an important role in infectious diseases and in autoimmune disorders ^18, 57–59^. We find anti-chemokine antibodies in these illnesses, but the patterns are different when compared to each other and to COVID-19. In HIV-1, a chronic viral infection, antibodies are significantly enhanced against more than half of all the chemokines, but do not include antibodies to either CCL19 or CXCL17, which are characteristic of COVID-19. Antibodies to the chemokine ligands of the HIV-1 coreceptors (CXCR4 and CCR5) are also detected at higher levels ^60^. In contrast, *Borrelia* infection does not induce development of anti-chemokine antibodies, with the sole exception of anti-CXCL14 IgGs. Autoantibodies in the three autoimmune disorders considered in our study (AS, RA, SjS) are generally similar to each other, but distinct from those in viral infection. We speculate that these differences reflect the unique role of chemokines in each of these diseases.

Infection can trigger antibody polyreactivity and autoimmunity that are generally deleterious ^61–63^. Since here we show that post-infectious autoantibodies can be associated with positive outcomes, we favor the view of post-infectious autoantibodies as disease modifiers. In COVID-19, the infection induces the expression of chemokines, leading to a pro-inflammatory milieu that clears infected cells but also causes collateral damage ^20, 24, 27–30^. Since anti-chemokine antibodies are present in plasma at concentrations able to modulate cellular migration, the variety and amount of anti-chemokine antibodies that are present or induced upon infection in each individual may modulate the quality and strength of the inflammatory response, which in turn would impact disease manifestation, severity and long COVID. This could also in part explain the variable lack of success of convalescent plasma treatment in COVID-19 ^64^, for which donors were selected based on virus neutralizing activity and not for the presence of autoantibodies that could modulate the inflammatory response.

We discovered and characterized the first, human-derived monoclonal antibodies against four chemokines, including anti-CXCL13 and anti-CXCL16 antibodies that are relevant as predictors of long COVID. Consistent with the 2-step model of chemokine receptor activation ^45, 65^, all the N-loop antibodies that were tested effectively reduced chemotaxis. Anti-inflammatory compounds, such as steroids and IL-6 blockers, are currently deployed in the clinic against COVID-19. Further studies are needed to determine whether agents that target the chemokine system could impact positively on the inflammatory phase of COVID-19 and reduce the development of long COVID.

## MATERIAL AND METHODS

### Major Resources Table

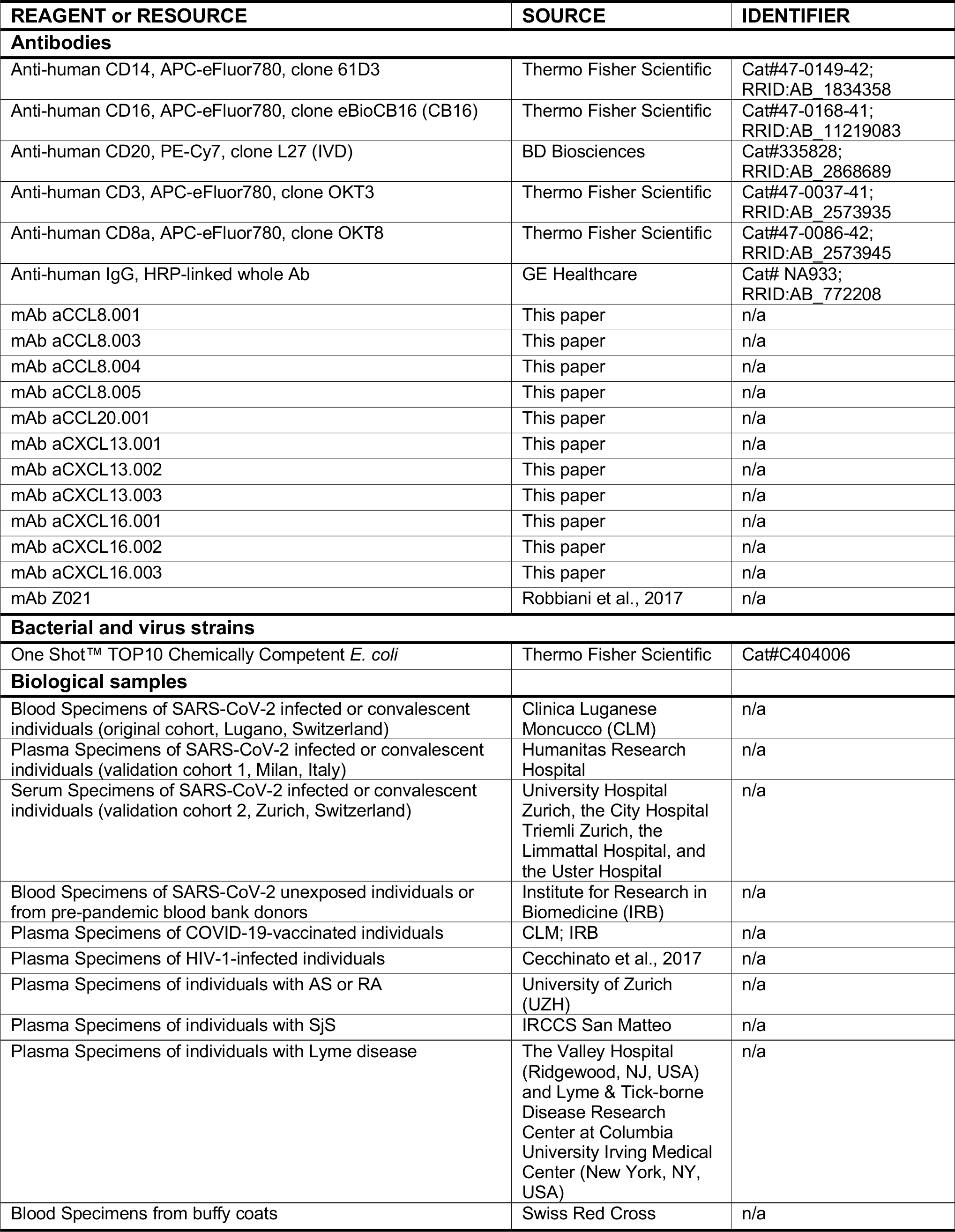
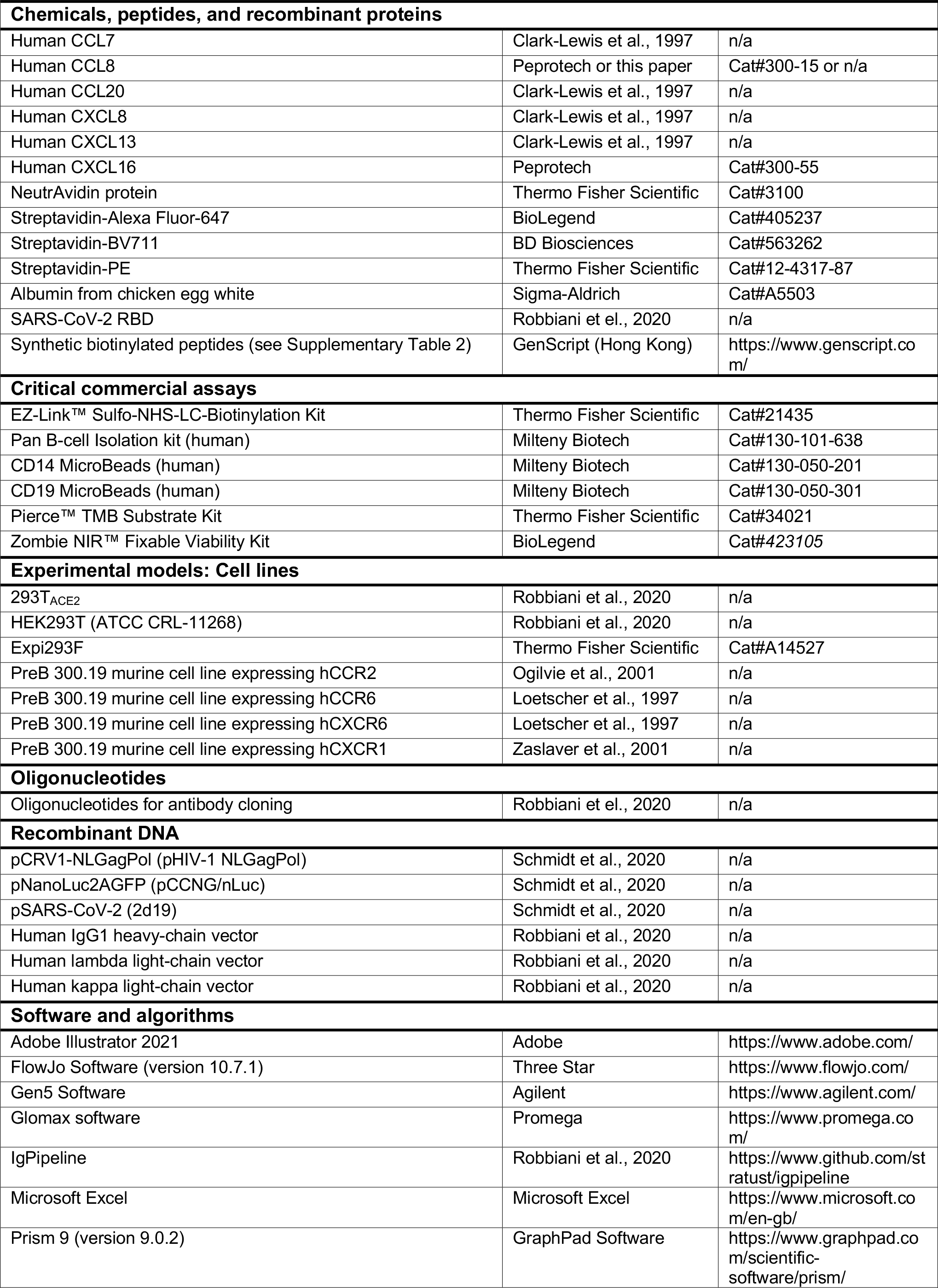
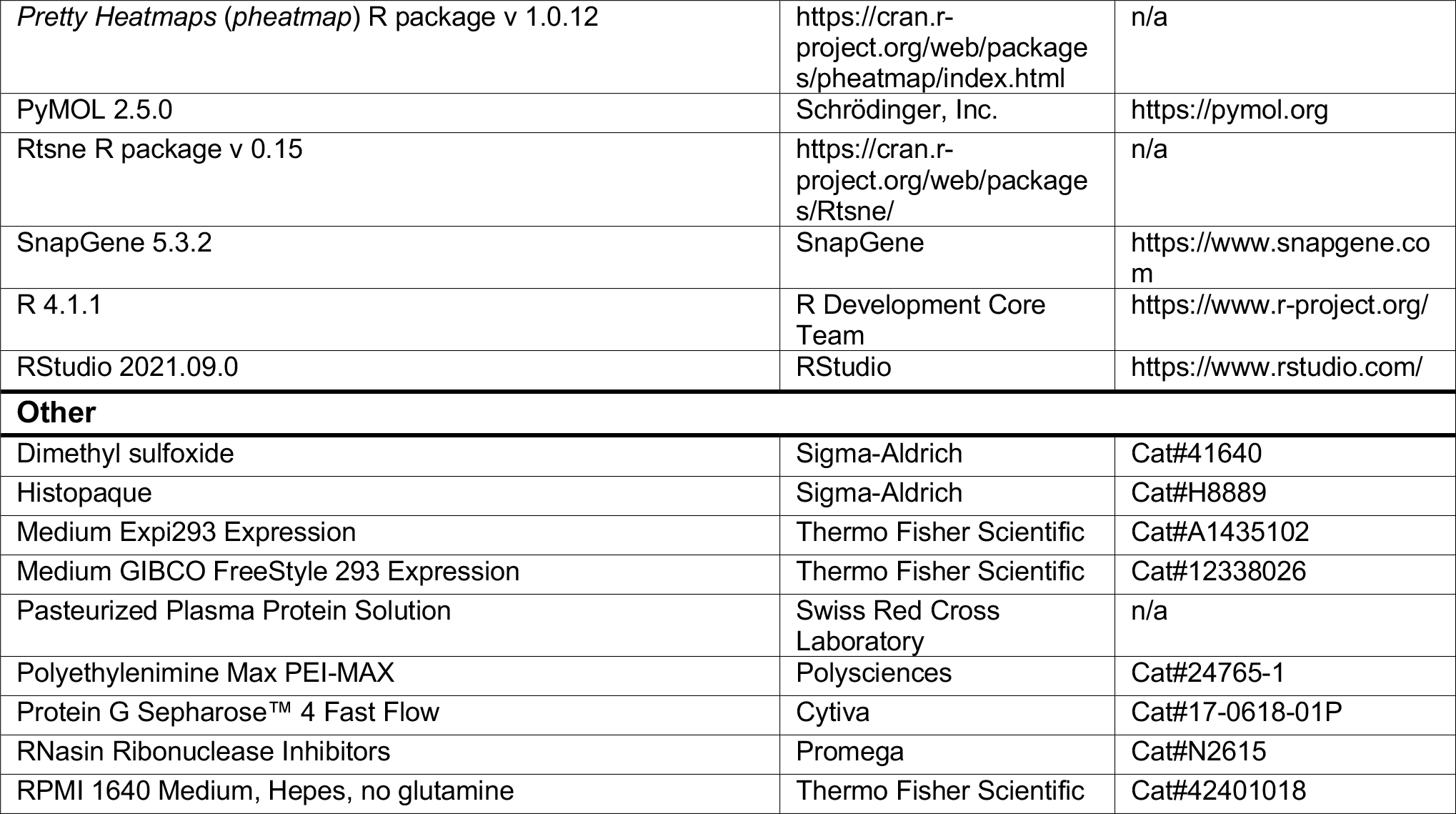

### Study participants and ethical approvals

#### COVID-19 original cohort (Lugano)

71 participants, diagnosed with COVID-19 at the Clinica Luganese Moncucco (CLM, Switzerland) between 08.03.2020 and 22.11.2020, were enrolled in the study and divided into two groups, according to the severity of the acute disease. The hospitalized group included 50 participants; the outpatient group included 21 close contacts of the hospitalized group, who only received at-home care. Inclusion criteria for the hospitalized group were a SARS-CoV-2 positive nasopharyngeal swab test by real-time reverse transcription-polymerase chain reaction (RT-PCR) and age ≥18 years. Inclusion criteria for the outpatient group were being a symptomatic close contact (living in the same household) of an individual enrolled in the hospitalized group and age ≥18 years. Serologic tests confirmed COVID-19 positivity for all the participants (Fig. 1a; Extended Data Fig. 4a). At the 12-month visit, participants were asked to indicate the presence or absence of persisting symptoms related to COVID-19 according to a questionnaire. The study was performed in compliance with all relevant ethical regulations and the study protocols were approved by the Ethical Committee of the Canton Ticino (ECCT): CE-3428 and CE-3960.

#### COVID-19 validation cohort 1 (Milan)

44 participants, diagnosed with COVID-19 and hospitalized at the Humanitas Research Hospital (Milan, Italy) between 10.03.2020 and 29.03.2021, were enrolled in the study. Inclusion criteria were a SARS-CoV-2 positive nasopharyngeal swab test by RT-PCR and age ≥18 years. Serologic tests confirmed COVID-19 positivity for the participants who were not tested by RT-PCR. Individuals were stratified in mild or severe depending on duration of hospitalization (mild: *≤*5 days; severe: *≥*7 days). The study was performed in compliance with all relevant ethical regulations and the study protocols were approved by the Ethical Committee of Humanitas Research Hospital (authorization n° 738/20 and n° 956/20).

COVID-19 validation cohort 2 (Zurich)^15^: 104 participants, diagnosed with COVID-19 at the University Hospital Zurich, the City Hospital Triemli Zurich, the Limmattal Hospital or the Uster Hospital between April 2020 and April 2021, were included in the study and divided into two groups, according to the severity of the acute disease. The hospitalized group included 38 participants, whereas the outpatient group included 66 individuals, who only received at-home care. Inclusion criteria for the participants were a SARS-CoV-2-positive nasopharyngeal swab test by RT-PCR and age ≥18 years. At the 13-month visit, blood was collected, and participants were asked by trained study physicians to indicate the presence or absence of persisting symptoms related to COVID-19. The study was performed in compliance with all relevant ethical regulations and the study protocols were approved by the Cantonal Ethics Committee of Zurich (BASEC #2016-01440).

#### Control cohort

15 adult participants (≥18 years) with self-reported absence of prior SARS-CoV-2 infection or vaccination (confirmed by negative serologic test, Extended Data Fig. 4a) were enrolled between November 2020 and June 2021. Additional 8 pre-pandemic samples were obtained from blood bank donors (ECCT: CE-3428). Serologic tests confirmed COVID-19 negativity for all controls (Fig. 1a; Extended Data Fig. 4a).

#### Vaccination cohort

16 adult participants (≥18 years) with self-reported absence of prior SARS-CoV-2 infection (confirmed by negative serologic test, Extended Data Fig. 5f) and who received two doses of mRNA-based COVID-19 vaccine ^66, 67^, were enrolled on the day of first vaccine dose or earlier, between November 2020 and June 2021 (ECCT: CE-3428).

#### HIV-1 and autoimmune diseases cohorts

Pre-pandemic plasma samples were obtained from the following participants: 24 HIV-1 positive (ECCT: CE-813) ^68^, 13 each with Ankylosing Spondylitis, Rheumatoid Arthritis (ECCT: CE-3065, and Ethical Committee of the Canton Zurich EK-515), or Sjögren’s syndrome (IRCCS Policlinico San Matteo Foundation Ethics Committee n.20070001302).

#### Lyme disease cohort

Plasma samples of 27 individuals with erythema migrans (Lyme disease) and 30 controls were obtained at The Valley Hospital (Ridgewood, NJ, USA) and Lyme & Tick-borne Disease Research Center at Columbia University Irving Medical Center (New York, NY, USA) between 2015 and 2019. All were between 18-89 years of age and all denied being immunocompromised. <u>Lyme disease cohort:</u> Individuals had new or recent onset erythema migrans, exposure to a Lyme endemic area in the prior 30 days and received no more than 3 weeks of antibiotic treatment. <u>Healthy control cohort:</u> Individuals reported being medically healthy, had an unremarkable physical exam and blood tests, had no signs or symptoms of infection or illness, denied having had a diagnosis and/or treatment for Lyme and/or another tick-borne disease within the past 5 years, and denied having a tick bite in the prior 6 months. The Lyme cohort samples were collected at the time of the erythema migrans and 6 months later in average. The study was performed in compliance with all relevant ethical regulations and the study protocol was approved by the New York State Psychiatric Institute Institutional Review Board (#6805).

Written informed consent was obtained from all participants, and all samples were coded. Demographic, clinical, and serological features of the cohorts are reported in Supplementary Tables 1 and 3.

### Blood collection, processing, and storage

Blood was collected by venipuncture at approximately 6-month intervals and the peripheral blood mononuclear cells (PBMCs) were isolated using Histopaque density centrifugation (Lugano and control cohort). Total PBMCs were aliquoted and frozen in liquid nitrogen in the presence of fetal calf serum and DMSO. Plasma was aliquoted and stored at -20°C or less. Before use, plasma aliquots were heat-inactivated (56°C for 1 h) and then stored at 4°C. For chemotaxis assays, CD14^+^ monocytes and CD19^+^ B cells were enriched from fresh PBMCs derived from blood donors (Swiss Red Cross Laboratory; ECCT: CE-3428) through positive immunoselection (130-050-201 and 130-050-301, respectively [Miltenyi Biotec, Bergisch Gladbach, Germany]) according to the manufacturer’s instructions. After isolation, CD19^+^ B cells were rested overnight in RPMI-1640 medium supplemented with 10% (v/v) fetal bovine serum (FBS), 1% (v/v) non-essential amino acids, 1 mM sodium pyruvate, 2 mM GlutaMAX, 50 μM β-Mercaptoethanol and 50 U/ml penicillin/streptomycin (all from Gibco) before being used in chemotactic assays. For the other cohorts, see references^15, 69^.

### Reagents

#### Peptides

Synthetic peptides containing the N-loop or the C-terminal sequence of human chemokines were designed and obtained (> 75% purity) from GenScript (Hong Kong). All peptides are biotinylated (biotin-Ahx) at the N-terminus and amidated at the C-terminus. In addition, the first 2-4 amino acids of each peptide (GS, GGS, GGGS, or GGK depending on the length of the N-loop/C-terminus of the chemokine) consist of a linker between the biotin and the chemokine sequence. Peptides are generally 25 amino acids long, or between 22-25 amino acids when synthesis was problematic. The sequence of the IFN*α*2 peptide (7-28) was based on a previously described immunoreactive epitope in myasthenia gravis patients ^70^, and the one from the SARS-CoV-2 nucleocapsid protein (N) peptide (157-178) was described in ^71^. An irrelevant peptide was used as negative control. The amino acid sequences of all peptides in this study are listed in Supplementary Table 2.

#### Proteins

CCL7, CCL20, CXCL8 and CXCL13 were synthesized using tBoc solid-phase chemistry ^72^. CCL8 and CXCL16 were obtained from Peprotech (Cat#300-15 and Cat#300-55, respectively) or produced and purified in house. Briefly, recombinant chemokines were expressed in *E. coli*, purified from inclusion bodies by immobilized-metal affinity chromatography, and folded under N2 protection in an arginine-containing buffer (80 mM Tris-Cl [pH 8.5], 100 mM NaCl, 0.8 M arginine, 2 mM EDTA, 1 mM cysteine, 0.2 mM cystine) as previously described ^73^. After recovery and concentration, the purification tag was cleaved with enterokinase, and the processed chemokine was purified by C18 reverse phase chromatography. The SARS-CoV-2 receptor binding domain (RBD) was produced and purified as described ^74^.

#### Chemotaxis

The migration of primary human monocytes and B cells isolated from buffy coats, or of murine preB 300.19 cells stably expressing the human chemokine receptors CCR2^75^, CCR6, CXCR1^76^ and CXCR6^77^ was assayed using 48-well Boyden chambers (Neuro Probe, Cabin John, MD) with polyvinylpyrrolidone-free polycarbonate membranes with pore size of 3 µm for primary human B cells and 5 µm for the other cell types, as previously described ^78^. Briefly, 10^5^ primary human B cells or 5x10^4^ primary human monocytes and murine preB 300.19 cells were diluted in RPMI-1640 supplemented with 20 mM Hepes, pH7.4, and 1% pasteurized plasma protein solution (5% PPL SRK; Swiss Red Cross Laboratory, Bern, Switzerland). Cells were then added to the upper wells and the chemokine (with or without antibodies) to the bottom wells. After 120 min of incubation for primary human B cells and 90 min for the other cell types, the membrane was removed, washed on the upper side with phosphate-buffered saline (PBS), fixed, and stained with DiffQuik. All assays were done in triplicate, and for each well the migrated cells were counted at 100-fold magnification in 5 randomly selected high-power fields (5HPF).

#### Inhibition of chemotaxis by monoclonal antibodies (Fig. 2f,h; Extended Data Fig. 8g,j)

Experiments were performed with monoclonal antibodies at a final concentration of 30 µg/ml (Extended Data Fig. 8g) or 50 µg/ml (Fig. 2f,h; Extended Data Fig. 8j). Baseline migration was determined in the absence of chemoattractant (buffer control).

#### Inhibition of chemotaxis by plasma purified IgGs (Extended Data Fig. 8k)

IgGs were purified from a subset of samples of the COVID-19 and uninfected control cohorts using Protein G Sepharose 4 Fast Flow (Cytiva) according to manufacturer’s instructions (plasma:resuspended beads at a 5:4 [v/v] ratio), buffer-exchanged and concentrated in PBS by Amicon Ultra-4 centrifugal filters (30 kDa cutoff, Millipore). Chemotaxis of preB 300.19 expressing CCR2 or CXCR1 was performed at a final IgG concentration of 200 µg/ml (IgG concentration in human serum: ∼10’000 µg/ml^49^), in the presence of the chemokine concentration resulting in peak migration when no antibodies were added (CCL7 [100nM], CCL8 [100nM], CXCL8 [1nM]).

### ELISA

To evaluate the antibodies’ binding to chemokine peptides, 96-well plates (ThermoFisher, 442404) or 384-well plates (ThermoFisher, 464718) were coated with 50 μl (or 10 μl for 384- well plates) per well of a 2μg/ml Neutravidin (Life Technologies, 31000) solution in PBS, overnight at room temperature. Plates were washed 4 times with washing buffer (PBS + 0.05% Tween-20 [Sigma-Aldrich]) and incubated with 50 μl (or 10 μl for 384-well plates) per well of a 50 nM biotinylated peptide solution in PBS for 1 h at room temperature. After washing 4 times with washing buffer, plates were incubated with 200 μl (or 50 μl for 384-well plates) per well of blocking buffer (PBS + 2% BSA + 0.05% Tween-20) for 2 h at room temperature.

Plates were then washed 4 times with washing buffer, and serial dilutions of monoclonal antibodies or plasma were added in PBS + 0.05% Tween-20 and incubated for 1 h at room temperature. To screen for the presence of anti-chemokine IgGs, plasma samples were assayed (unless otherwise stated) at 1:50 starting dilution followed by 3 fourfold serial dilutions (1:200, 1:800, 1:3200). Monoclonal antibodies were tested at 5 μg/ml starting concentration followed by 11 threefold serial dilutions. Plates were subsequently washed 4 times with washing buffer and incubated with anti-human IgG secondary antibody conjugated to horseradish peroxidase (HRP) (GE Healthcare, NA933) at a 1:5000 dilution in PBS + 0.05% Tween-20. Finally, after washing 4 times with washing buffer, plates were developed by the addition of 50 μl (or 10 μl for 384-well plates) per well of the HRP substrate TMB (ThermoFisher, 34021) for 10 min. The developing reaction was stopped with 50 μl (or 10 μl for 384-well plates) per well of a 1 M H2SO4 solution, and absorbance was measured at 450 nm with an ELISA microplate reader (BioTek) with Gen5 software. A positive control (broadly reactive plasma from donor CLM70) and negative control (uninfected participant) samples were included in each experiment. Since the basal average optical density likely also depends on intrinsic features of each peptide that is used to coat the ELISA plate, the presented values should be interpreted as relative rather than absolute. The Area Under the Curve (AUC) was obtained from two independent experiments and plotted with GraphPad Prism. The main findings were further confirmed by assaying subsets of samples belonging to the different groups, side-by-side on the same plates (data not shown).

#### Lyme disease cohort (Extended Data Fig. 9b)

Plasma was assayed at a 1:100 starting dilution, followed by 2 additional four-fold dilutions (1:400 and 1:1600).

#### Reactivity at 6 versus 12 months (Extended Data Fig. 5b,c)

Experiments were performed with plasma samples from different time points side-by-side on the same plate. In Extended Data Fig. 5b, plasma was assayed at a 1:50 starting dilution, followed by 4 additional fivefold dilutions. Anti-RBD IgG levels were measured in COVID-19 convalescents, who had not received a COVID-19-mRNA vaccine between first and second visit (no vaccination) or in individuals with at least one dose of vaccine at least 10 days before blood sampling at the second visit (Extended Data Fig. 5b; see Supplementary Table 1).

#### Kinetic of signature anti-chemokine IgG antibodies (Extended Data Fig. 5e)

Experiments were performed with plasma samples from different time points assayed at 1:50 dilution side-by-side on the same plate, and the average optical density at 450 nm obtained from two independent experiments was plotted with GraphPad Prism.

#### IgG antibodies binding to SARS-CoV-2 RBD (Figs. 1a and 2b; Extended Data Fig. 4a,c,d; Extended Data Fig. 5b,f; Extended Data Fig. 6c)

Experiments were performed with 96-well plates coated with 50 μl per well of a 5 μg/ml protein solution in PBS overnight at room temperature, and subsequently blocked and treated as described above. In this case, plasma samples were assayed either at a 1:50 starting dilution followed by 7 additional threefold serial dilutions (Figs. 1a and 2b; Extended Data Fig. 4a,c,d; Extended Data Fig. 6c) or followed by 3 additional fivefold serial dilutions (Extended Data Fig. 5b,f).

#### Chemokine quantification in plasma

Plasma levels of 14 chemokines were measured using the Luminex Discovery Assay - Human Premixed Multi-Analyte Kit (R&D Systems, LXSAHM-14) following the manufacturer’s instructions. Chemokines included in the panel were: CCL2, CCL3, CCL4, CCL19, CCL21, CCL22, CCL25, CXCL2, CXCL5, CXCL8, CXCL9, CXCL10, CXCL13 and CXCL16. Each sample was measured in duplicate using a Luminex FLEXMAP 3D system.

#### Single cell sorting by flow cytometry

B cells were enriched from PBMCs of uninfected controls or of COVID-19 convalescent individuals 6 months after COVID-19 (participant CLM9 for anti-CCL8 antibodies; CLM64 for anti-CCL20 antibodies; CLM5, CLM7 and CLM33 for anti-CXCL13 antibodies; and CLM8 and CLM30 for anti-CXCL16 antibodies), using the pan-B-cell isolation kit according to manufacturer’s instructions (Miltenyi Biotec, 130-101-638). The enriched B cells were subsequently stained in FACS buffer (PBS + 2% FCS + 1mM EDTA) with the following antibodies/reagents (all 1:200 diluted) for 30 min on ice: anti-CD20-PE-Cy7 (BD Biosciences, 335828), anti-CD14-APC-eFluor 780 (Thermo Fischer Scientific, 47-0149-42), anti-CD16- APC-eFluor 780 (Thermo Fischer Scientific, 47-0168-41), anti-CD3-APC-eFluor 780 (Thermo Fischer Scientific, 47-0037-41), anti-CD8-APC-eFluor 780 (Invitrogen, 47-0086-42), Zombie NIR (BioLegend, 423105), as well as fluorophore-labeled ovalbumin (Ova) and N-loop peptides. Live single Zombie-NIR^−^CD14^−^CD16^−^CD3^−^CD8^−^CD20^+^Ova^−^N-loop-PE^+^N-loop- AF647^+^ B cells were single-cell sorted into 96-well plates containing 4 μl of lysis buffer (0.5× PBS, 10 mM DTT, 3,000 units/ml RNasin Ribonuclease Inhibitors [Promega, N2615]) per well using a FACS Aria III, and the analysis was performed with FlowJo software. The anti-CCL20 antibody sequences were obtained by sorting with a pool of 12-peptides; for all the others, a single peptide was used. The sorted cells were frozen on dry ice and stored at −80 °C.

#### Antibody sequencing, cloning, production and purification

Antibody genes were sequenced, cloned and expressed as previously reported ^79–81^. Briefly, reverse-transcription of RNA from FACS-sorted single cells was performed to obtain cDNA, which was then used for amplification of the immunoglobulin IGH, IGK and IGL genes by nested PCR. Amplicons from this first PCR reaction served as templates for sequence and ligation independent cloning (SLIC) into human IgG1 antibody expression vectors. Monoclonal antibodies were produced by transiently transfecting Expi293F cells cultured in Freestyle-293 Expression Medium (ThermoFisher) with equal amounts of Ig heavy and light chain expression vectors using polyethylenimine Max (PEI-MAX, Polysciences) as a transfection reagent. After 6-7 days of culture, cell supernatants were filtered through 0.22 μm Millex-GP filters (Merck Millipore), and antibodies were purified using Protein G Sepharose 4 Fast Flow (Cytiva) according to manufacturer’s instructions and buffer-exchanged and concentrated in PBS by Amicon Ultra-4 centrifugal filters (30 kDa cutoff, Millipore). Where indicated, the anti-Zika virus monoclonal antibody Z021 ^79^ was used as an isotype control.

#### Computational analysis of antibody sequences

Antibody sequences were analyzed using a collection of Perl and R scripts provided by IgPipeline and publicly available on GitHub (https://github.com/stratust/igpipeline) ^46^. In brief, sequences where annotated using IgBlast ^82^ v 1.14.0 with IMGT domain delineation system and the Change-O toolkit v 0.4.5 ^83^. CDR3 sequences were determined by aligning the IGHV and IGLV nucleotide sequence against their closest germlines using the blastn function of IgBlast.

#### SARS-CoV-2 pseudotyped reporter virus and neutralization assay

To generate (HIV-1/NanoLuc2AEGFP)-SARS-CoV-2 particles, HEK293T cells were co-transfected with the three plasmids pHIVNLGagPol, pCCNanoLuc2AEGFP, and SARS-CoV-2 S as described elsewhere ^46, 84^. Supernatants containing virions were collected 48 h after transfection, and virion infectivity was determined by titration on 293TACE2 cells. The plasma neutralizing activity was measured as previously reported ^46, 84^.

Briefly, threefold serially diluted plasma samples (from 1:50 to 1:328’050) were incubated with SARS-CoV-2 pseudotyped virus for 1h at 37 °C, and the virus-plasma mixture was subsequently incubated with 293TACE2 cells for 48 h. Cells were then washed with PBS and lysed with Luciferase Cell Culture Lysis 5× reagent (Promega). Nanoluc Luciferase activity in cell lysates was measured using the Nano-Glo Luciferase Assay System (Promega) with Modulus II Microplate Reader User interface (TURNER BioSystems). The obtained relative luminescence units were normalized to those derived from cells infected with SARS-CoV-2 pseudotyped virus in the absence of plasma. The NT50 values were determined using four-parameter nonlinear regression with bottom and top constrains equal to 0 and 1, respectively (GraphPad Prism). The dotted line (NT50=5) in the plots represents the lower limit of detection of the assay.

#### Model interaction between chemokine and chemokine receptor

The illustrative model in Extended Data Fig. 1b was generated from the structure of inactive CCR2 (PDB code: 5T1A) ^85^, together with the electron microscopy structures of CCR5 and CCR6 (PDB codes: 6MEO and 6WWZ, respectively) ^86, 87^ by using SWISS-MODEL ^88^ server and the molecular graphics program PyMOL 2.5.0 for modeling the N- and C-terminus of the receptor. The crystal structure of CCL8 (MCP-2) (PDB code: 1ESR) ^89^, and the electron microscopy structure of CCR6 ^87^ were used to model the complex. The intracellular residues were removed for clarity.

### Statistical analysis

#### Tests for statistical significance

Upon testing of parametric assumptions, statistical significance between two groups was determined using either parametric paired two-tailed Student’s t test, or non-parametric two-tailed Mann–Whitney U-tests (unpaired samples), or Wilcoxon signed-rank test (paired samples). Statistical significance between more than two groups was evaluated using Kruskal-Wallis test (followed by Dunn multiple comparisons), one-way ANOVA (followed by Tukey multiple comparisons), or two-way Repeated Measures ANOVA (followed by Šídák multiple comparisons), as described in the figure legends. Statistical significance of the signature chemokines (CCL19, CCL22, CXCL17, CXCL8, CCL25, CXCL5, CCL21, CXCL13 and CXCL16) was also confirmed when applying the Bonferroni criterion in order to guarantee a familywise level of significance equal to 0.05. Statistical significance from a 2x2 contingency table was determined with Fisher’s exact test.

Correlations were assessed using Pearson correlation analysis. A p-value of less than 0.05 was considered statistically significant. In the figures, significance is shown as follow: ns p≥0.05 (not significant), *p<0.05, **p<0.01, ***p<0.001 and ****p<0.0001. Data and statistical analyses were performed with GraphPad Prism.

#### t-SNE

t-SNE analysis was performed using the Rtsne R package v 0.15 (https://CRAN.R-project.org/package=Rtsne) using the AUC values for all chemokines. The theta parameter for the accuracy of the mapping was set to zero in all cases for exact t-SNE.

#### Clustering

Hierarchical clustering was created using the hclust R function v 4.1.1. Clustering analysis was performed using the correlation as distance and the Ward’s method as agglomerative criterion. Heatmaps were created with either GraphPad Prism (Fig. 1a; Extended Data Fig. 2d) or the *Pretty Heatmaps* (*pheatmap*) R package v 1.0.12 (Extended Data Fig. 10b). In Extended Data Fig. 10b, each column containing a distinct chemokine was scaled with the scaling function provided by R, which sets the mean and the standard deviation to 0 and 1, respectively.

#### Logistic regression and additional analyses

Logistic regression was performed using the GLM (Generalized Linear Models) function provided by the R package v 4.1.1. To identify which variables to include in the analysis, AUCs were ranked according to the p-value obtained with a Mann-Whitney-Wilcoxon nonparametric test on the Lugano cohort. The first N variables minimizing the AIC (Akaike information criterion) were then used in the fitting. Furthermore, the same set of variables was used to perform the fitting with the Milan and Zurich cohorts. In each plot, values from 0 to 0.5 and from 0.5 to 1 on the y-axis represent the assignment of individuals to the A and B groups (of a Prediction A versus B; see grey backgrounds), respectively. On the x-axis, samples are divided into the two groups and subsequently ordered according to sample ID as shown in Supplementary Table 1. Dots in the grey area represent individuals that are assigned to the correct group. We additionally performed χ^2^-tests considering covariates that are known to influence COVID-19 severity (demographics [gender and age] and comorbidities [diabetes and cardiovascular diseases]) and found that none of them was significantly different between groups. Race/ethnicity was not analyzed because the cohort is nearly 100% Caucasian; similarly, immune deficiency was rare and was not considered. Logistic regression analysis using the combination of these covariates (age, gender, diabetes and cardiovascular diseases) allowed proper assignment with accuracies of 74.6% (COVID-19 severity; outpatient vs hospitalized) and 68.3% (Long COVID; no Sx vs *≥*1 Sx). Notably, the accuracy using anti-chemokine antibody values is even better (77.5% [COVID-19 severity]) and 77.8% [Long COVID]). These analyses are shown in Supplementary Table 7.

## Acknowledgments

We thank all study participants and their families, the medical personnel of the Clinica Luganese Moncucco; Carol Sotsky, Nancy Rioux, Shreya Doshi, and Tarek Hijazi (Lyme disease cohort); Marina Sironi, Roberto Leone, Monica Rimoldi and the Humanitas COVID- 19 Task Force (Milan cohort); Sara Hasler, Sarah Adamo, Miro Raeber, Jakob Nilsson, Esther Bächli, Alain Rudiger and Lars Huber (Zurich cohort) and Sandra Jovic (Institute for Research in Biomedicine) for their support and technical assistance. We also acknowledge Thiago Oliveira (Rockefeller University) for sharing the script for the clustering analysis, and Theodora Hatziioannou and Paul Bieniasz (Rockefeller University) for sharing plasmids and protocols for SARS-CoV-2 pseudovirus. ACa thanks Mrs. Flora Gruner for the generous support. This study was also in part financed within the framework of the Swiss HIV Cohort Study using data gathered by the Five Swiss University Hospitals, two Cantonal Hospitals, 15 affiliated hospitals and 36 private physicians (listed in http://www.shcs.ch/180-health-care-providers). Members of the Swiss HIV Cohort Study are listed in https://shcs.ch/184-for-shcs-publications. We are grateful to Marco Baggiolini for mentoring, support and fruitful discussions.

## Funding

Swiss Vaccine Research Institute (SVRI) (DFR)

National Institutes of Health grant U01 AI151698 (United World Antiviral Research Network, UWARN) (DFR)

National Institutes of Health grant P01 AI138938 (DFR)

National Institutes of Health grant U19 AI111825 (DFR)

European Union’s Horizon 2020 research and innovation programme grant 101003650 (DFR, LV)

Fidinam Foundation (MUg)

Rocca Foundation (Mug, AMant)

Ceschina Foundation (MUg)

Swiss HIV Cohort Study (#719), Swiss National Science Foundation (#201369), and SHCS Research Foundation (MUg, EB, AR)

Italian Ministry of Health for COVID-19 COVID-2020-12371640 (AMant) Dolce & Gabbana fashion house (AMant)

Swiss National Science Foundation: NRP 78 Implementation Programme (CC, OB), #4078P0-198431 (OB), #310030-200669 (OB), and #310030-212240 (OB)

Clinical Research Priority Program CYTIMM-Z of University of Zurich (UZH) (OB)

Pandemic Fund of UZH (OB)

Innovation grant of USZ (OB)

Digitalization Initiative of the Zurich Higher Education Institutions Rapid-Action Call #2021.1_RAC_ID_34 (CC)

Swiss Academy of Medical Sciences (SAMW) fellowships #323530-191220 (CC) and #323530-191230 (YZ)

Lyme & Tick-Borne Diseases Research Center at Columbia University Irving Medical Center, established by the Global Lyme Alliance, Inc and the Lyme Disease Association, Inc. (BAF)

## Author contributions

Conceptualization: JMu, VCe, ACa, MUg, DFR

Software: MM, SM, JS, ACa

Validation: JMu, VCe, ACa

Formal Analysis: JMu, VCe, ACa, SM, MBLR

Investigation: JMu, VCe, AAS, EG, JMo, TG, FBi

Resources: JMu, TG, PP, FBi, VCr, LP, CT, GD-S, BM, MT, DJ, ST, MP, LV, MB, PAM, AF-P, CG, MUh, EB, AR, ACi, AManz, RF, FBa, AV, GDN, CC, PT, YZ, LAM, BB, AMant, BAF, OB

Writing – Original Draft: JMu, VCe, MUg, DFR Writing – Review & Editing: JMu, VCe, MUg, DFR Supervision: DFR, MUg

Funding Acquisition: DFR, MUg, ACa, LV, AR, EB, BAF, OB, CC, AMant

## Data and materials availability

All data are available in the main text or the supplementary materials. Computer code for antibody sequence, logistic regression, clustering and t-SNE analyses will be deposited at GitHub upon publication (https://github.com). Material is available upon request and may require MTA.

## Extended Data Figures

**Extended Data Fig. 1.**
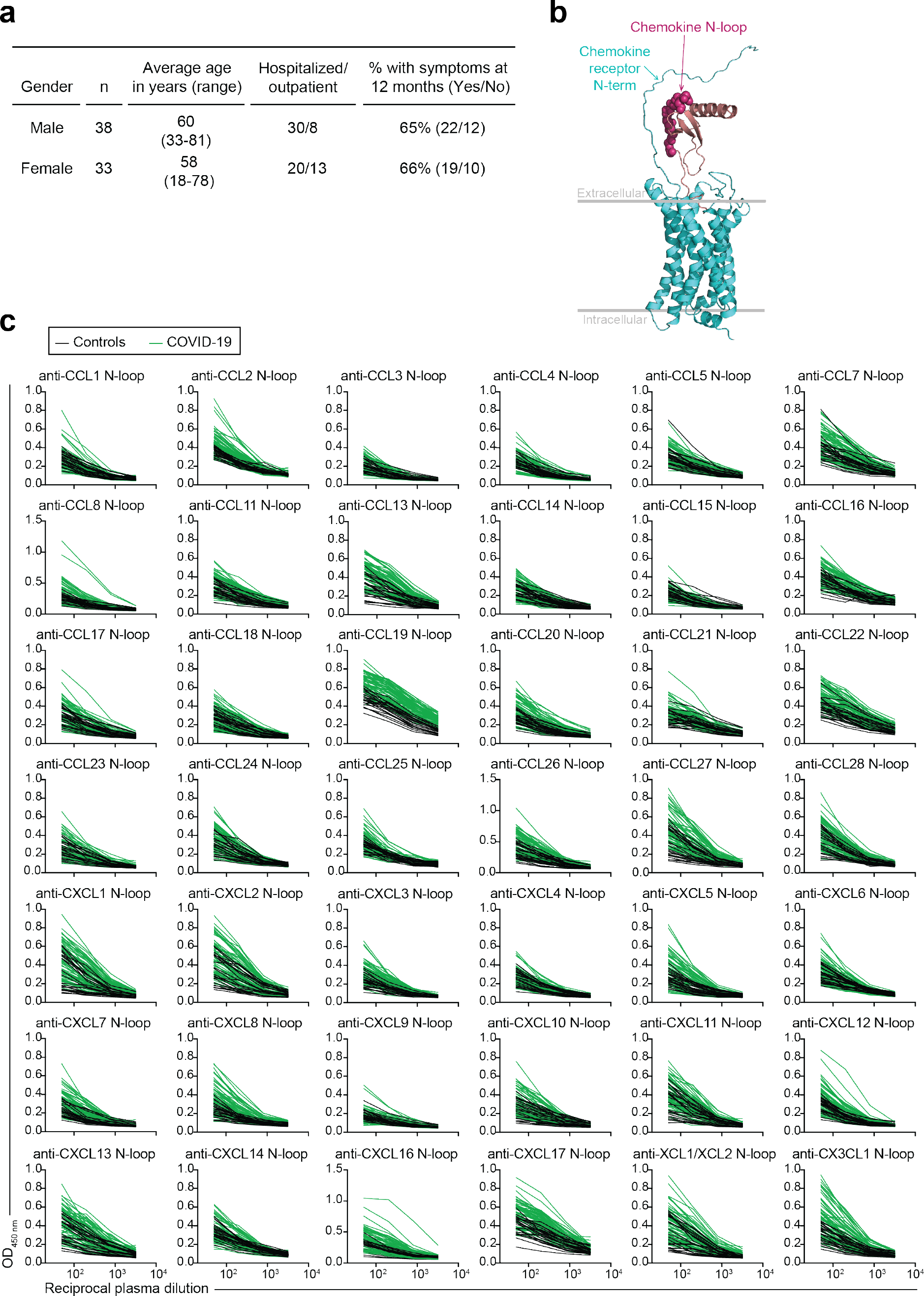
Anti-chemokine N-loop antibodies in COVID-19, related to Fig. 1. (a) Characteristics of the Lugano COVID-19 cohort. (**b**) Model of the interaction between a chemokine and its receptor. Arrows point to the area of putative interaction between the N-terminus of the receptor and the chemokine N-loop (shown by spheres). Chemokine is magenta and chemokine receptor is cyan. (**c**) The amount of plasma IgG antibodies against each chemokine N-loop was determined by ELISA for COVID-19 convalescents (n=71) and controls (n=23). Average optical density (OD450) measurements of two independent experiments.

**Extended Data Fig. 2.**
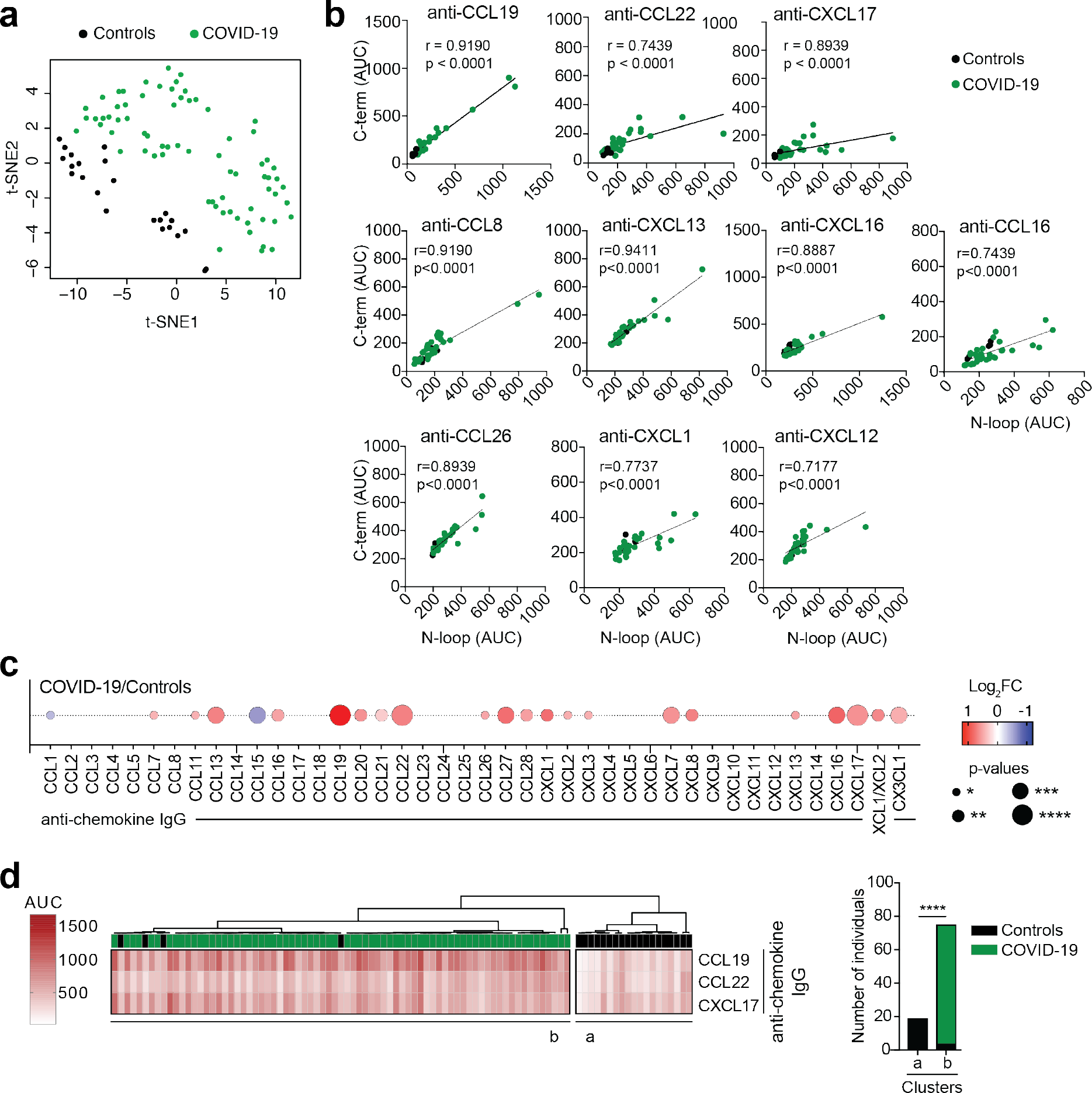
Analyses of anti-chemokine antibodies, related to Fig. 1. **(a)** t-SNE distribution of COVID-19 convalescents and controls, as determined with the 42 datasets combined. (**b**) Pearson correlations of antibodies to the N-loop and C-terminal peptides of the same chemokine. ELISA was performed in a cohort subset (Controls, n=5; COVID-19, n=31). Average of two independent experiments. (**c**) Differences in anti-chemokine antibodies between groups. Summary circle plot: circle size indicates significance; colors show the Log2 fold-change increase (red) or decrease (blue) in the COVID-19 group over control. Two-tailed Mann–Whitney U-tests. (**d**) Antibodies to CCL19, CCL22 and CXCL17 classify COVID-19 convalescents versus controls. Unsupervised hierarchical clustering analysis with the COVID- 19 signature antibodies. The distribution of the groups within each cluster is also shown. Fisher’s exact test.

**Extended Data Fig. 3.**
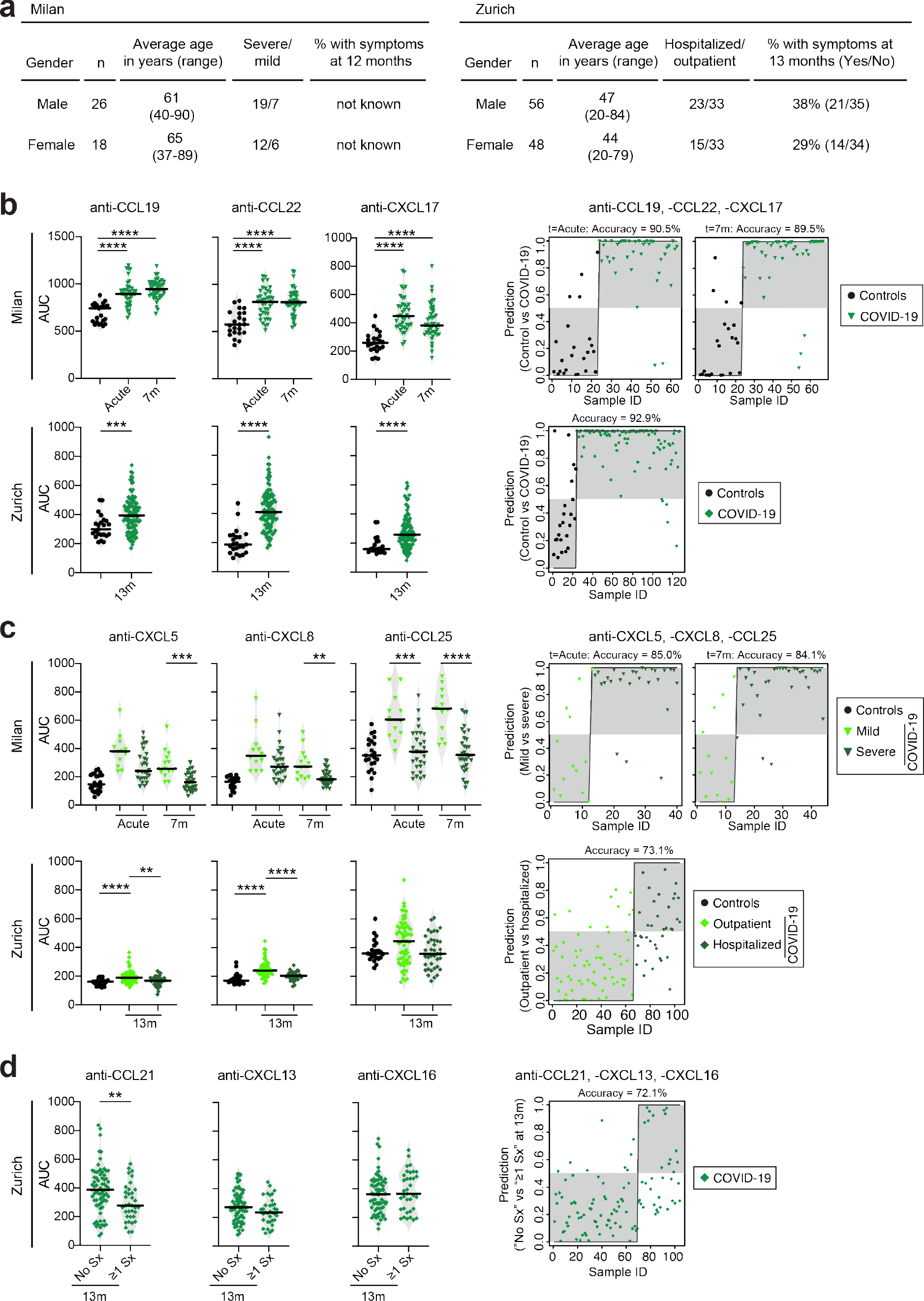
Analyses of anti-chemokine antibodies in the validation cohorts of Milan and Zurich, related to Figs. 1-3. **(a)** Characteristics of the COVID-19 cohorts. (**b**) Left, difference in autoantibodies to CCL19, CCL22 and CXCL17 (COVID-19 signature). Kruskal-Wallis test followed by Dunn’s multiple comparison test (Milan) and two-tailed Mann– Whitney U-tests (Zurich). Right, assignment of COVID-19 convalescents and controls based on the COVID-19 signature antibodies by logistic regression analysis. (**c**) Left, difference in autoantibodies to CXCL5, CXCL8 and CCL25 (COVID-19 hospitalization signature). Kruskal-Wallis test followed by Dunn’s multiple comparison test. Right, assignment of COVID-19 mild/outpatient and severe/hospitalized individuals based on the COVID-19 hospitalization signature antibodies by logistic regression analysis. (**d**) Left, difference in autoantibodies to CCL21, CXCL13 and CXCL16 (Long COVID signature). Two-tailed Mann– Whitney U-tests. Right, group assignment based on the Long COVID signature antibodies by logistic regression analysis in the Zurich cohort. In (**b-d**), horizontal bars indicate median values; dots on grey background are correctly assigned; data are shown as average AUC of two independent experiments.

**Extended Data Fig. 4.**
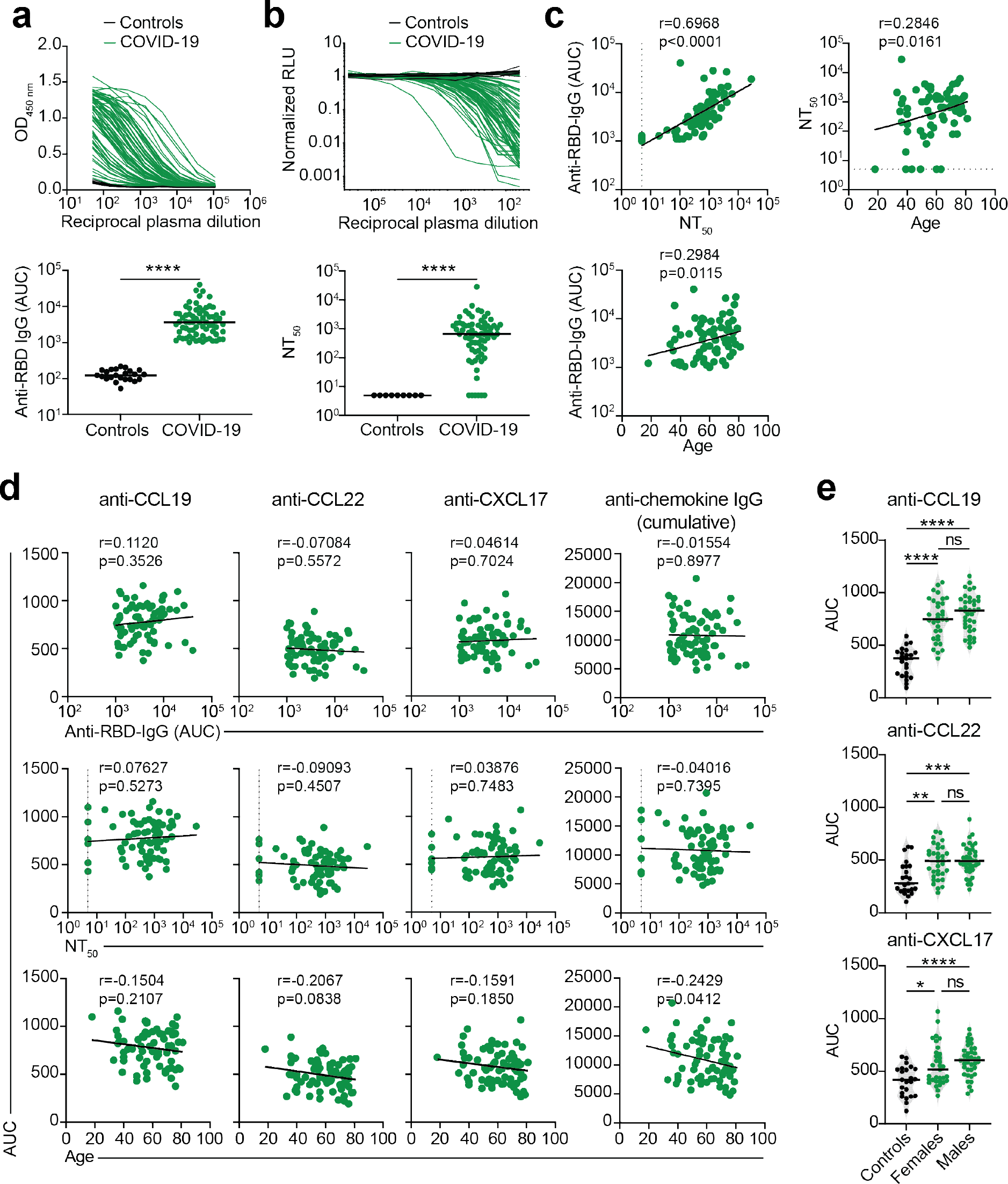
Correlation analyses of COVID-19 signature anti-chemokine antibodies, related to Fig. 1. (**a**) Anti-RBD IgG antibodies in the cohort samples. Top, optical density reactivity (OD450) of serial plasma dilutions to the receptor binding domain (RBD) of SARS-CoV-2 Spike, as determined by ELISA. Bottom, AUC of the data in the top panel. COVID-19 convalescents (n=71); controls (n=23). Average of two independent experiments. Horizontal bars indicate median values. Two-tailed Mann–Whitney U-tests. (**b**) Plasma neutralizing activity against SARS-CoV-2 pseudovirus. Top, relative luciferase units (RLU) normalized to no plasma control. Bottom, half-maximal neutralizing titers (NT50) based on the data in the top panel. COVID-19 convalescents (n=71); controls (n=9). Average of two independent experiments. Horizontal bars indicate median values. Two-tailed Mann–Whitney U-tests. (**c**) Pearson correlations of anti-RBD IgG and NT50 values to each other and with age. Average of two independent experiments. (**d**) Pearson correlations of anti-chemokine IgG with anti-RBD IgG, NT50 values and age. COVID-19 signature antibodies individually, and cumulative signal of the IgGs against the peptides for all 43 chemokines. (**e**) Analysis of anti- signature chemokines IgG by gender. Data are shown as average AUC of two independent experiments. Horizontal bars indicate median values. Kruskal-Wallis test followed by Dunn’s multiple comparison test.

**Extended Data Fig. 5.**
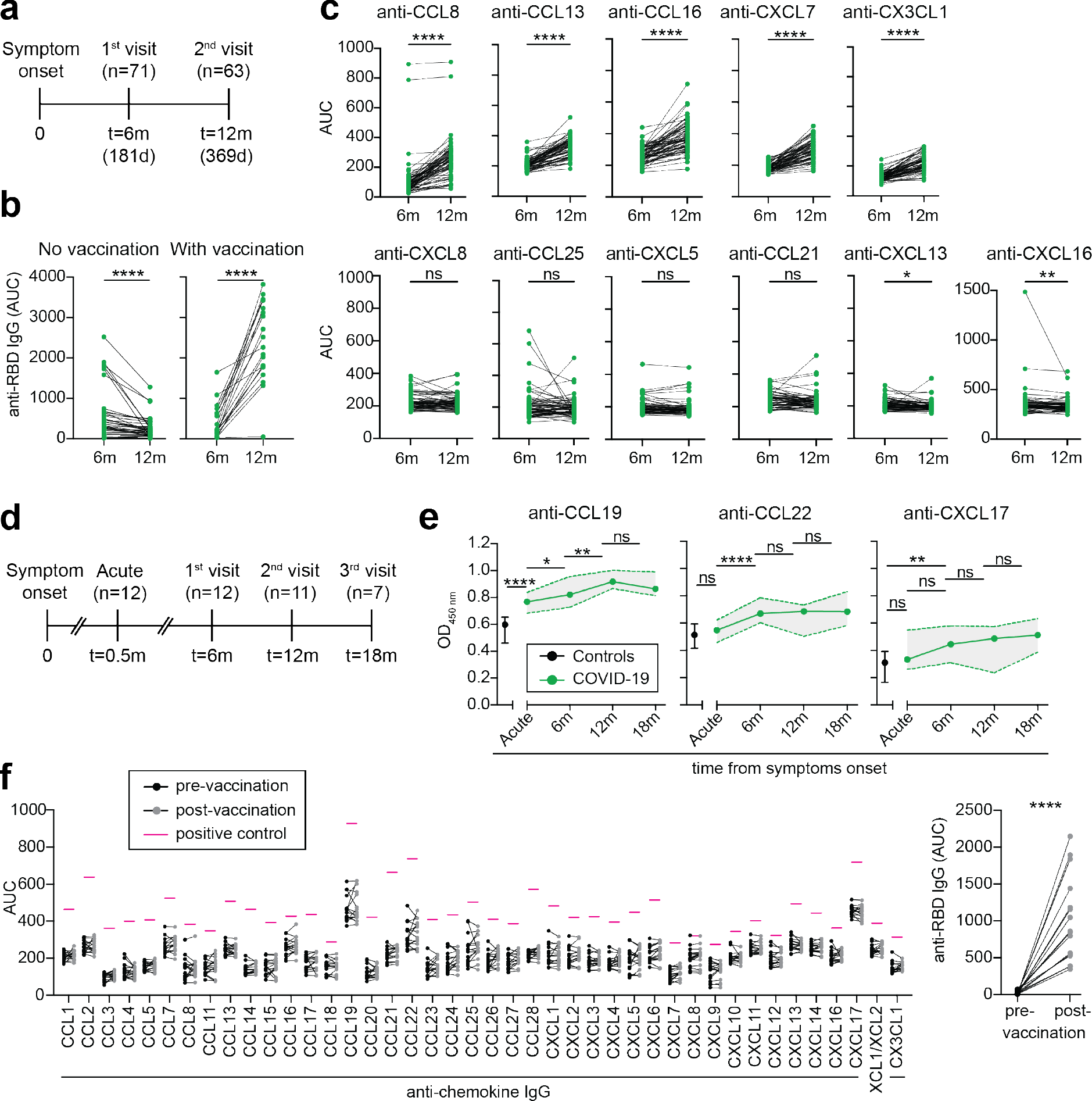
Anti-chemokine antibodies over time and upon COVID-19 vaccination, related to Fig. 1. (**a**) Diagram of the time points of blood collection after onset of COVID-19 symptoms. (**b**) Anti-RBD IgG antibodies at 6 and 12 months in vaccinated and non-vaccinated convalescents, as determined by ELISA. Average AUC from two independent experiments. Wilcoxon signed-rank test. (**c**) Anti-chemokine IgG antibodies at 6 and 12 months in convalescents. AUC from two independent experiments. Wilcoxon signed-rank test. (**d**) Diagram of the time points of blood collection after onset of COVID-19 symptoms in a subset of COVID-19 hospitalized individuals. (**e**) Anti-chemokine IgG antibodies at 15 days (Acute), 6, 12 and 18 months after onset of COVID-19 symptoms. Average optical density (OD450) values from two independent experiments. One-way ANOVA test followed by Tukey’s multiple comparison test. Data are shown as median±range. (**f**) Anti-chemokine IgG antibodies before and approximately 4 months after COVID-19 mRNA vaccination of uninfected individuals (n=16). AUC from two independent experiments. Pink lines represent the signal of a positive control plasma sample with broad reactivity (CLM70). Anti-RBD IgG is shown alongside as control (right panel). Wilcoxon signed-rank test with false discovery rate (FDR) approach.

**Extended Data Fig. 6.**
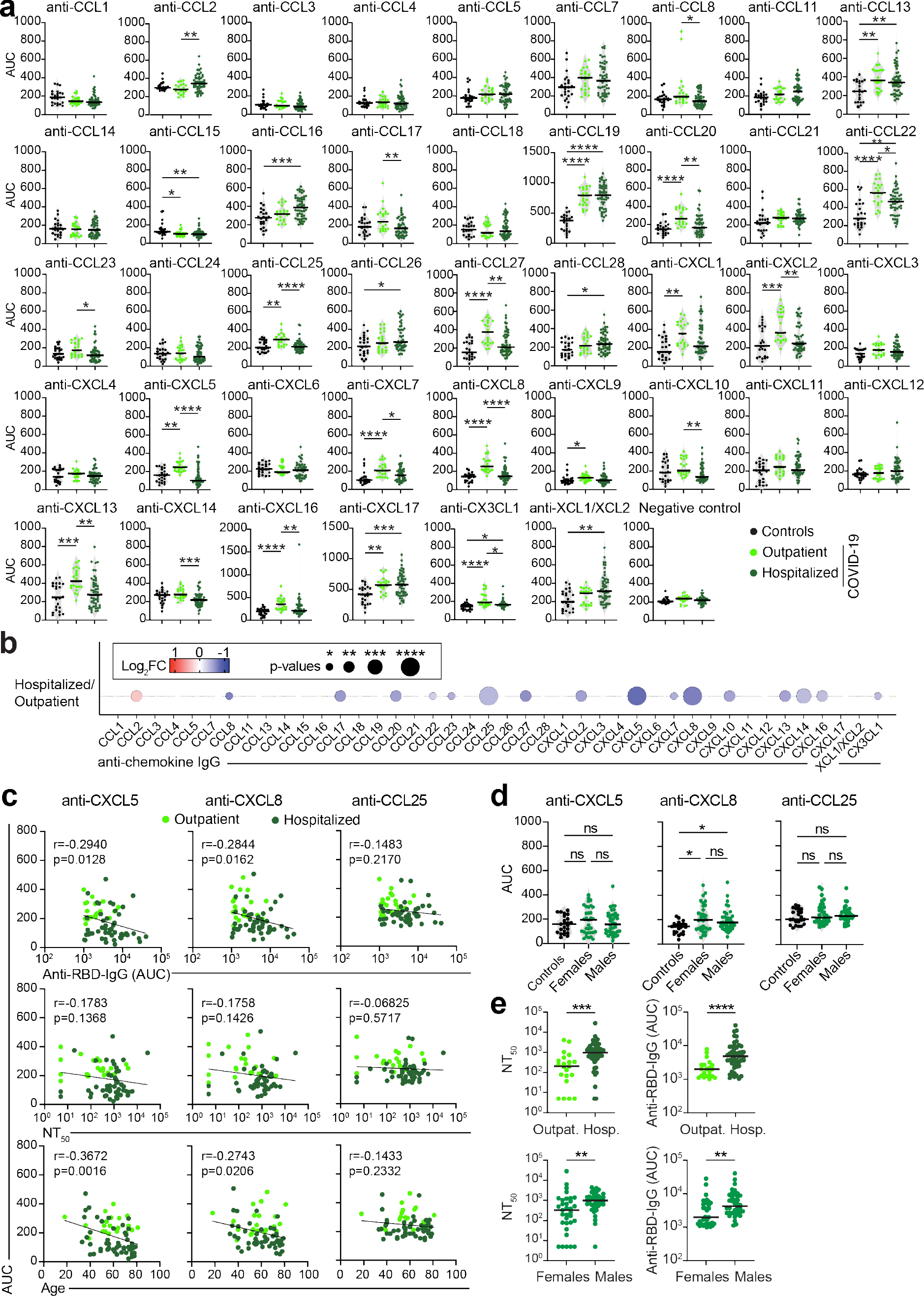
Anti-chemokine antibodies in COVID-19 outpatient and hospitalized individuals and correlation analyses of COVID-19 hospitalization signature antibodies, related to Fig. 1. (**a**) Difference in anti-chemokine antibodies in outpatient versus hospitalized individuals at 6 months. Average AUC of two independent experiments. Horizontal bars indicate median values. Kruskal-Wallis test followed by Dunn’s multiple comparison test. (**b**) Difference in anti-chemokine antibodies between COVID-19 outpatient and hospitalized individuals. Summary circle plot: circle size indicates significance; colors show the Log2 fold-change increase (red) or decrease (blue) over outpatients. Kruskal-Wallis test followed by Dunn’s multiple comparison test. (**c**) Pearson correlations of anti-chemokine IgGs with anti-RBD IgG, NT50 values and age. COVID-19 hospitalization signature antibodies individually. Average of two independent experiments. (**d**) Analysis of anti- signature chemokine IgGs by gender. Data are shown as average AUC of two independent experiments. Horizontal bars indicate median values. Kruskal-Wallis test followed by Dunn’s multiple comparison test. (**e**) Analysis of anti-RBD IgG and NT50 values by group and by gender. Average of two independent experiments. Horizontal bars indicate median values. Two-tailed Mann–Whitney U-tests.

**Extended Data Fig. 7.**
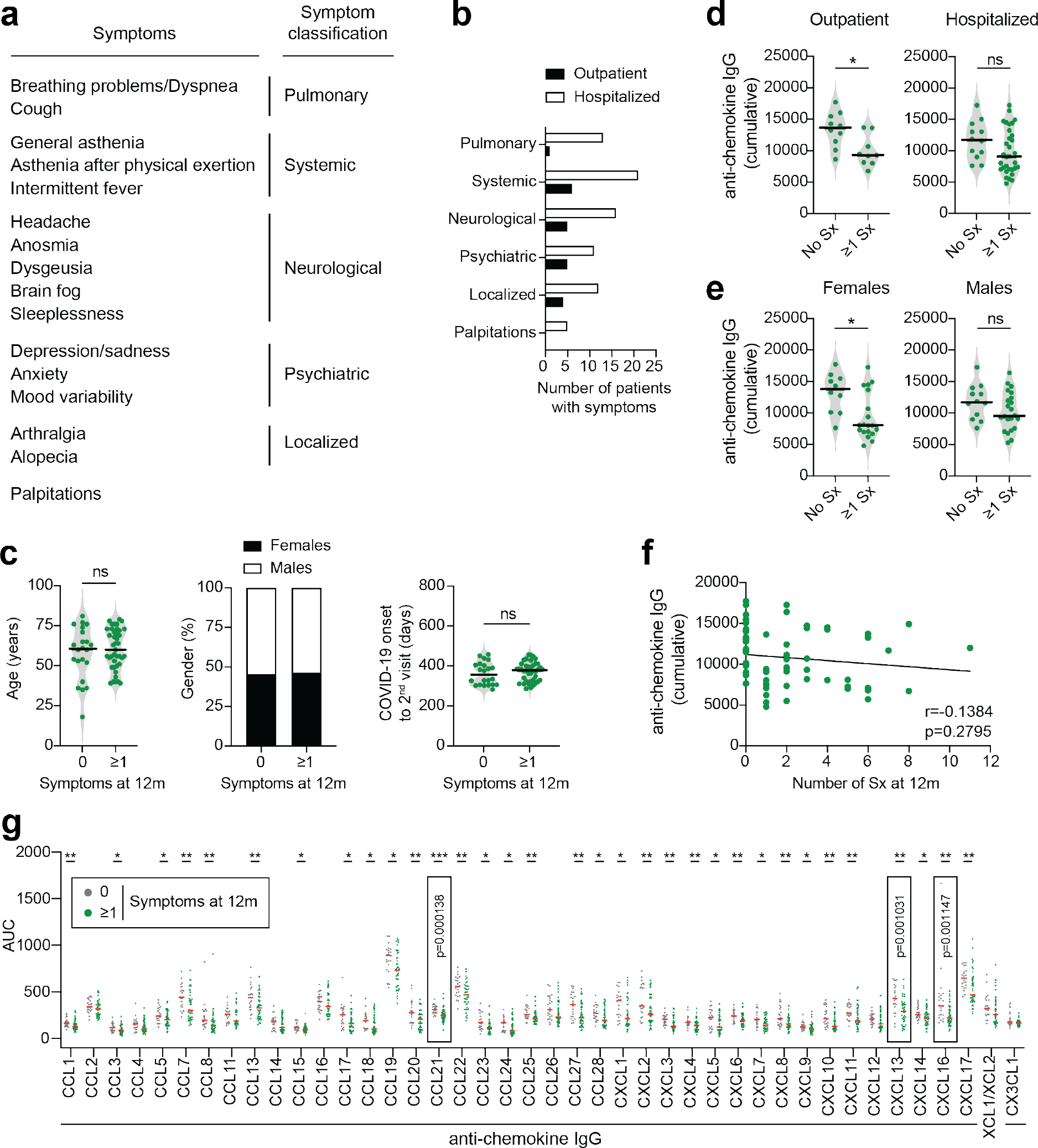
Anti-chemokine antibodies and long-term COVID-19 symptoms, related to Fig. 2. (**a**) Classification of long-term COVID-19 symptoms at 12 months (t=12m). Incidence of symptoms at 12 months. Participants are grouped in outpatient and hospitalized individuals. (**c**) Analysis of age (left), gender distribution (middle) and time from COVID-19 onset to 2^nd^ visit (t=12m; right). Horizontal bars indicate median values. Two-tailed Mann–Whitney U-tests. (**d,e**) Difference in cumulative anti-chemokine antibodies according to the presence or absence of symptoms at 12 months in disease severity groups (d) or by gender (e). Data are shown as average AUC of two independent experiments. Horizontal bars indicate median values. Two-tailed Mann–Whitney U-tests. (**f**) Pearson correlation of anti-chemokine antibodies and the number of symptoms at 12 months. Average of two independent experiments. (**g**) Difference in anti-chemokines antibodies at 6 months and the presence or absence of symptoms at 12 months. Data are shown as average AUC of two independent experiments. Horizontal bars indicate median values. Two-tailed Mann–Whitney U-tests. The exact P-value is given for the 3 chemokines displaying the highest significance.

**Extended Data Fig. 8.**
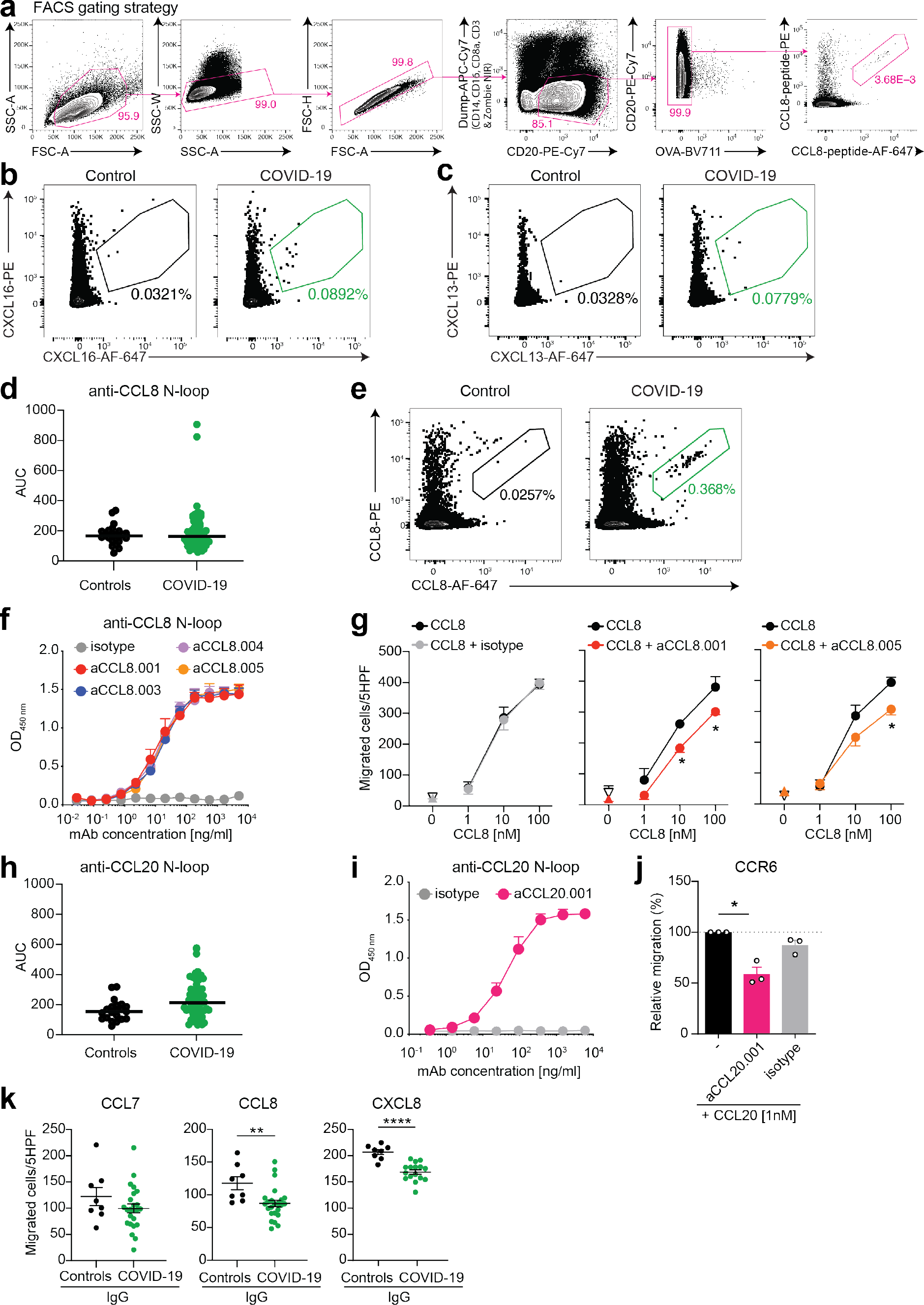
Human monoclonal antibodies that impede chemotaxis, related to Fig. 2. (**a**) Gating strategy for sorting CCL8 N-loop specific B cells by flow cytometry. (**b**) Human B cells specific for CXCL16. Representative flow cytometry plots identifying human B cells binding to the CXCL16 N-loop peptide (gate). The frequency of antigen-specific B cells is shown. (**c**) Human B cells specific for CXCL13. Representative flow cytometry plots identifying human B cells binding to the CXCL13 N-loop peptide (gate). The frequency of antigen-specific B cells is shown. (**d**) Identification of individuals with high anti-CCL8 N-loop antibodies. Area under the curve (AUC), as determined by ELISA. Average of two independent experiments. COVID-19 convalescents (n=71); controls (n=23). Horizontal bars indicate median values. (**e**) CCL8 binding human B cells. Flow cytometry plots identify human B cells binding to the CCL8 N-loop peptide (gate). The frequency of antigen-specific B cells is shown. (**f**) Monoclonal antibodies to the CCL8 N-loop. ELISA binding curves of representative antibodies. Average of two independent experiments (Mean+SEM). (**g**) Chemotaxis of human monocytes towards CCL8 is inhibited by monoclonal antibodies. Mean±SEM of migrated cells in 5 high-power fields (HPF). At least 3 independent experiments with cells from different donors. Up-pointing triangle is antibody alone, and down-pointing triangle is buffer control. Two-way RM ANOVA followed by Šídák’s multiple comparisons test. (**h**) Identification of individuals with high anti-CCL20 N-loop antibodies. Area under the curve (AUC), as determined by ELISA. Average of two independent experiments. COVID-19 convalescents (n=71); controls (n=23). Horizontal bars indicate median values. (**i**) Monoclonal antibodies to the CCL20 N-loop. ELISA binding curve of a representative antibody. Average of two independent experiments (Mean+SEM). (**j**) Anti-CCL20 N-loop antibodies inhibit CCL20 chemotaxis to CCR6. Relative cell migration towards CCL20. Mean+SEM of at least 3 independent experiments. Two-tailed Mann–Whitney U-tests. (**k**) Polyclonal IgGs from COVID-19 convalescents inhibit chemotaxis. Chemotaxis of preB 300.19 cells expressing either CCR2 or CXCR1 towards the indicated chemokines (CCL7 or CCL8 for CCR2; CXCL8 for CXCR1) was measured in the presence of plasma IgGs from COVID-19 convalescents (n=24 for CCL7 and CCL8; n=16 for CXCL8) or controls (n=8). Technical triplicates (Mean±SEM) of migrated cells in 5 high-power fields (HPF). Two-tailed Mann–Whitney U-tests.

**Extended Data Fig. 9.**
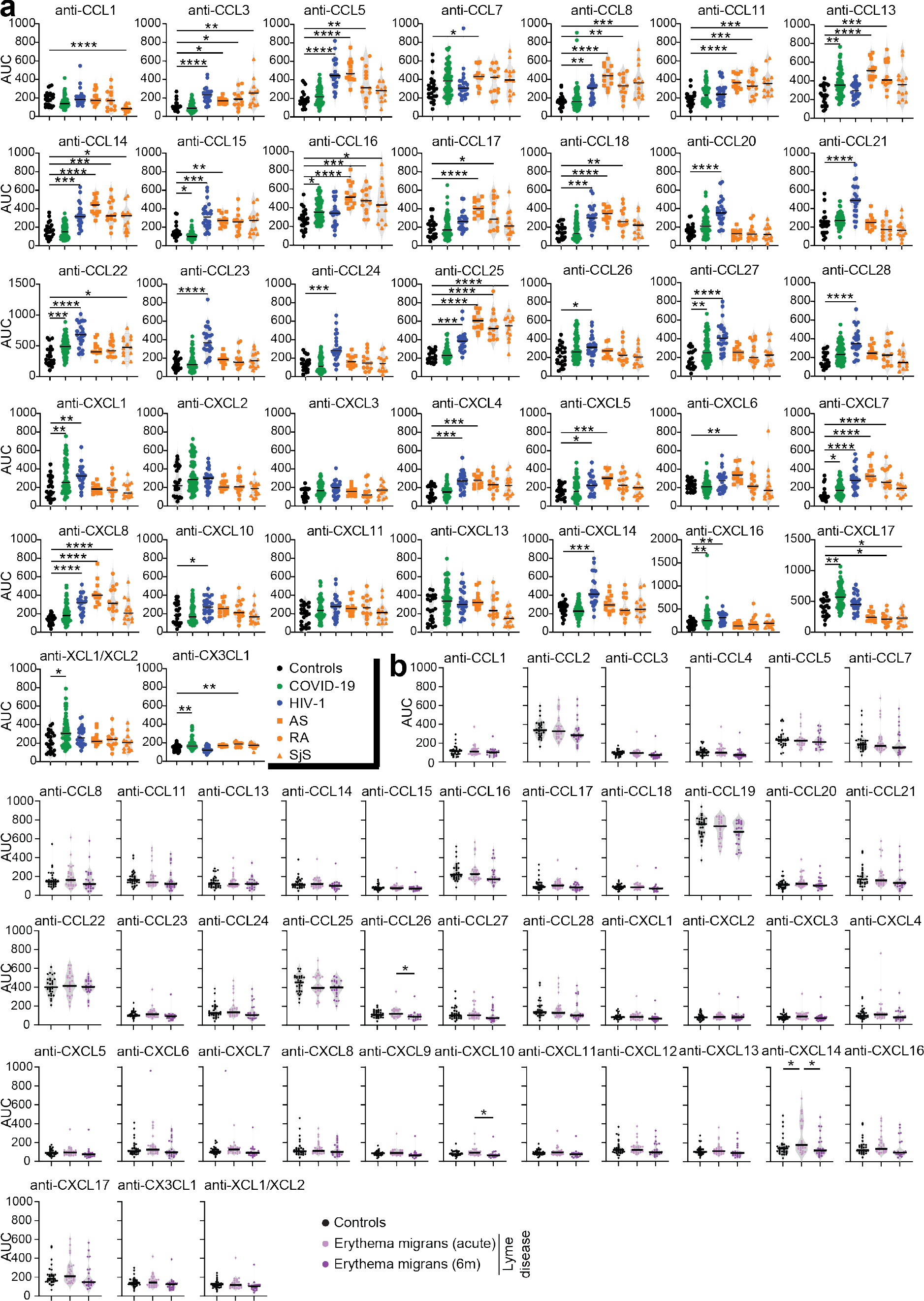
Anti-chemokine N-loop antibodies in HIV-1, autoimmune and Lyme diseases, related to Fig. 4. (**a**) The amount of plasma IgG antibodies against each chemokine N-loop was determined by ELISA for HIV-1 infected (n=24, blue) and autoimmune patients (n=39, orange). Autoimmune patients were subdivided in Ankylosing Spondylitis (AS, n=13), Rheumatoid Arthritis (RA, n=13), and Sjögren’s syndrome (SjS, n=13). Values from controls (n=23, black), and COVID-19 convalescents (n=71, green) are shown alongside. Average AUC of two independent experiments. Horizontal bars indicate median values. Statistical significance was determined using Kruskal-Wallis test followed by Dunn’s multiple comparison test over rank of the control group. (**b**) The amount of plasma IgG antibodies against each chemokine N-loop was determined by ELISA for *Borrelia*-infected patients (n=27, Lyme disease with erythema migrans) in the acute phase and at 6 months post infection, as measured by ELISA. Average AUC of two independent experiments. Horizontal bars indicate median values. Statistical significance was determined using Kruskal-Wallis test followed by Dunn’s multiple comparison test.

**Extended Data Fig. 10.**
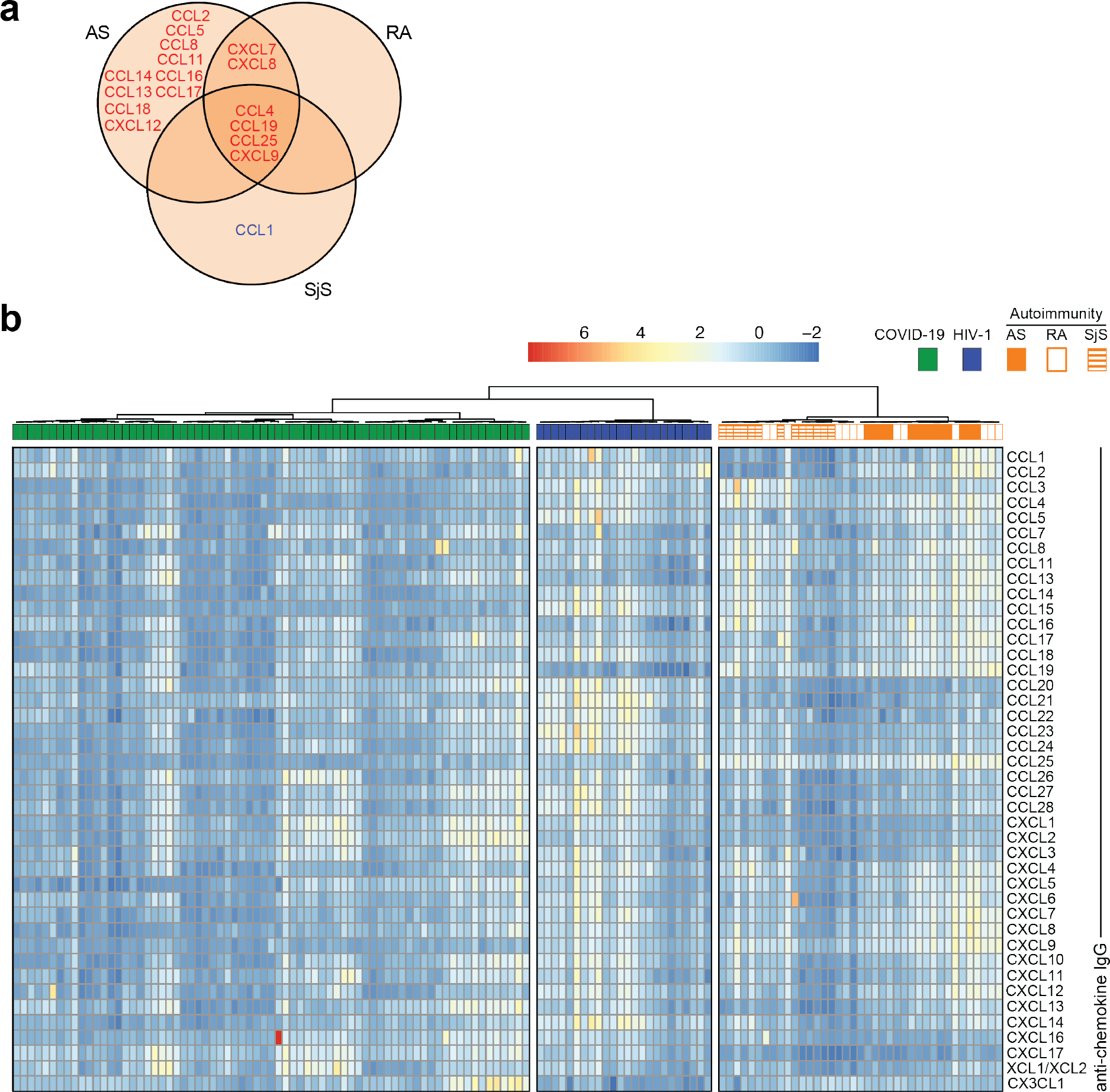
Clustering of COVID-19, HIV-1 and autoimmune diseases based on anti-chemokine antibodies, related to Fig. 4. (**a**) Venn diagram of the chemokines targeted by autoantibodies across the autoimmune disorders AS, RA and SjS. Red and blue colors indicate either an increase or decrease over controls with p<10^-4^. (**b**) Anti-chemokine antibodies correctly classify COVID-19 convalescents, HIV-1-infected, and patients with autoimmune disorders. Heatmap representing the normalized plasma IgG binding to 42 peptides comprising the N-loop sequence of all 43 human chemokines. Unsupervised hierarchical clustering analysis. The distribution of the groups within each cluster is shown.

